# A Thin Film Transistor Backplane for Scalable Chronic Neural Interfaces

**DOI:** 10.64898/2026.06.23.733868

**Authors:** Andrew M. Bourhis, Ritwik Vatsyayan, Karen J. Tonsfeldt, Ian Galton, Shadi A. Dayeh

**Affiliations:** Department of Electrical and Computer Engineering, University of California San Diego, La Jolla, CA, 92093 USA; Department of Obstetrics, Gynecology, and Reproductive Sciences, Center for Reproductive Science and Medicine, University of California San Diego, La Jolla, California 92093, United States; Department of Bioengineering, Department of Nanoengineering, Department of Neurological Surgery, and Materials Science and Engineering Program, University of California San Diego, La Jolla, California 92093, United States

## Abstract

Scaling neural interfaces to ever-higher channel counts has accelerated rapidly with advances in thin-film fabrication, lithography, and connectorization, enabling passive arrays to reach thousands of channels and chart credible pathways to much larger formats. Integrating active electronics directly at the sensing sites offers a complementary route to higher channel density by reducing the number of interconnects required to access large arrays. Here we introduce a monolithic flexible thin-film integrated circuit platform for active neural sensing, inspired by active-matrix display technology. The system integrates dual-gate amorphous indium gallium zinc oxide transistors on polyimide substrates to implement in-pixel transconductance amplification and row-column time-division multiplexing, improving scability for high-channel-count applications. Co-optimization of device architecture, contact engineering, and a hybrid ceramic-polymer thin-film encapsulation yields stable operation with projected lifetimes exceeding 38 years under accelerated aging. In acute and chronic *in vivo* rat studies, the platform exhibits negligible thermal burden, robust sensory-evoked recordings, and stable functionality over 30 days despite tissue encapsulation. These results establish display-inspired flexible thin-film electronics as a scalable building block for next-generation neural interfaces.

## Introduction

Over the past century, advances in neurotechnology have produced tools capable of recording neural activity across a wide range of spatial and temporal scales. Yet despite this progress, building systems capable of sampling the full tapestry of neural activity remains a major challenge. Electrocorticography (ECoG) and related surface-based interfaces capture the integrated activity of large neuronal populations across the cortex, offering a unique balance between signal stability, coverage, and invasiveness^1^. Progress in this space has consistently required navigating trade-offs among spatial resolution, cortical coverage, and surgical burden from early low-density surface electrodes to modern microfabricated arrays capable of dense sampling over increasingly large cortical areas^2–6^.

Recent advances in thin-film fabrication, lithography, and connectorization have enabled passive high-density electrode arrays with an order-of-magnitude higher channel count than the best prior designs, and credible pathways now exist to scale passive systems further than previously thought possible^4–7^. Such scaling, however, introduces engineering challenges, including increased connector width, interconnect routing density, and multi-layer metallization complexity, as each additional channel requires a dedicated electrical lead. Thin film technologies have enabled continued scaling by reducing interconnect pitch footprint, but this comes at the cost of increased fabrication complexity and mechanical considerations that must be carefully managed for chronic implantation and clinical translation.

Active electrode arrays, which integrate local transistors in the same substrate as the sensing electrodes, provide a complementary strategy for scaling neural interfaces by decreasing the number of external interconnects required to access large arrays. Silicon-based implementations of such systems have demonstrated promising electrical performance, including low noise, high power efficiency, and sophisticated on-chip signal processing capabilities, and have played a central role in advancing high-channel-count neural recording technologies ^8–13^. For chronic applications, however, the mechanical mismatch between rigid silicon substrates and soft neural tissue introduces challenges related to long-term reliability and tissue response. Approaches aimed at improving mechanical compliance such as aggressive substrate thinning, narrow probe geometries, or transfer of silicon devices onto flexible carriers ^3,14–16^ have shown promise but can also introduce additional considerations, including increased fracture susceptibility, overreliance on brittle inorganic encapsulation layers, and added constraints on long-term hermiticity in physiological environments. More fundamentally, the electronic properties of silicon are intrinsically linked to its crystalline bulk structure, which limits the extent to which mechanical compliance can be engineered independently of device performance.

Thin-film transistor (TFT) technologies, which are widely deployed in active-matrix organic light-emitting diode (AMOLED) displays (including most modern smartphones and smartwatches)^17,18^, offer an alternative electronic platform on which transistor device function is largely decoupled from the mechanical properties of the underlying substrate. TFTs can be fabricated directly on polymeric and other flexible materials using low-temperature processes, enabling substrate-agnostic circuit integration. The downsides are that the amorphous or polycrystalline nature of thin-film semiconductors result in reduced carrier mobility, elevated noise and off-state current leakage, and long-term electrical instability compared to crystalline silicon^19–22^. In addition, the monolithic integration of complementary TFTs remains an active area of research^23^, placing greater demands on the performance and stability of monopolar semiconductor technologies in practical circuit implementations. As a result, many prior active neural interfaces have relied on silicon-based transistors with post-fabrication mechanical adaptations or heterogeneous integration schemes^24,25^, which can increase process complexity and impose additional challenges for scalable, clinically robust systems.

In this work, we present a fully monolithic flexible integrated circuit (FIC) platform for active neural sensing, inspired by active-matrix display architectures. The platform employs dual-gate amorphous indium gallium zinc oxide (a-IGZO) TFTs fabricated directly on polyimide substrates and encapsulated using a multilayer atomic-layer-deposited aluminum oxide thin-film barrier. Through systematic co-optimization of contact engineering, device architecture, and encapsulation strategy, we realize a 256-channel electrocorticography array implementing row-column addressing with in-pixel transconductance amplification. This architecture reduces interconnect scaling from linear proportionality to square root proportionality while preserving mechanical compliance and signal fidelity comparable to passive surface arrays.

The platform was validated through extensive benchtop characterization, including over one year of accelerated aging tests projecting operational lifetimes exceeding 38 years, followed by acute and chronic *in vivo* experiments in Sprague-Dawley rats. Acute sensory-evoked recordings demonstrated negligible thermal burden^26–32^ and robust signal quality relative to a matched passive array. A 30-day chronic implantation confirmed stable electrical operation, maintained encapsulation integrity^33–40^, and captured consistent spatiotemporal cortical signals despite the expected development of fibrotic tissue observed upon explant. This work represents the first demonstration of a fully monolithic flexible TFT-based active neural interface maintaining functionality over multiple weeks *in vivo*, and establishes design principles for translating display-inspired thin-film electronics to scalable implantable neural interfaces.

## Results and Discussion

### Flexible Thin Film Integrated Circuit Platform for Neural Sensing

Inspired by active-matrix organic light emitting diode (AMOLED) display technology, we developed a monolithic thin film transistor fabrication process enabling fully integrated flexible neural interfaces on biocompatible polyimide substrates (Fig. 1). Our platform employs dual-gate amorphous indium gallium zinc oxide (a-IGZO) TFTs, selected for their exceptional electron mobility (∼10-20 cm^2^/Vs), low-temperature processing compatibility (deposited at <=200°C), and substrate-agnostic fabrication. Unlike typical silicon-based approaches, which require post-fabrication transfer or aggressive thinning, the fabrication process (Supplementary Fig. 1) produces inherently flexible circuits without compromising hermetic encapsulation (Supplementary Fig. 2-5) or introducing mechanical failure modes associated with brittle semiconductors. Furthermore, the monolithic fabrication process allows for greater design flexibility compared to heterogeneous integration strategies which may, for example, rely on thick inorganic layers to mitigate seam or void formation around transferred layers. A monolithic fabrication strategy on the other hand can achieve much finer feature size and superior morphological feature control.

**Figure 1.**
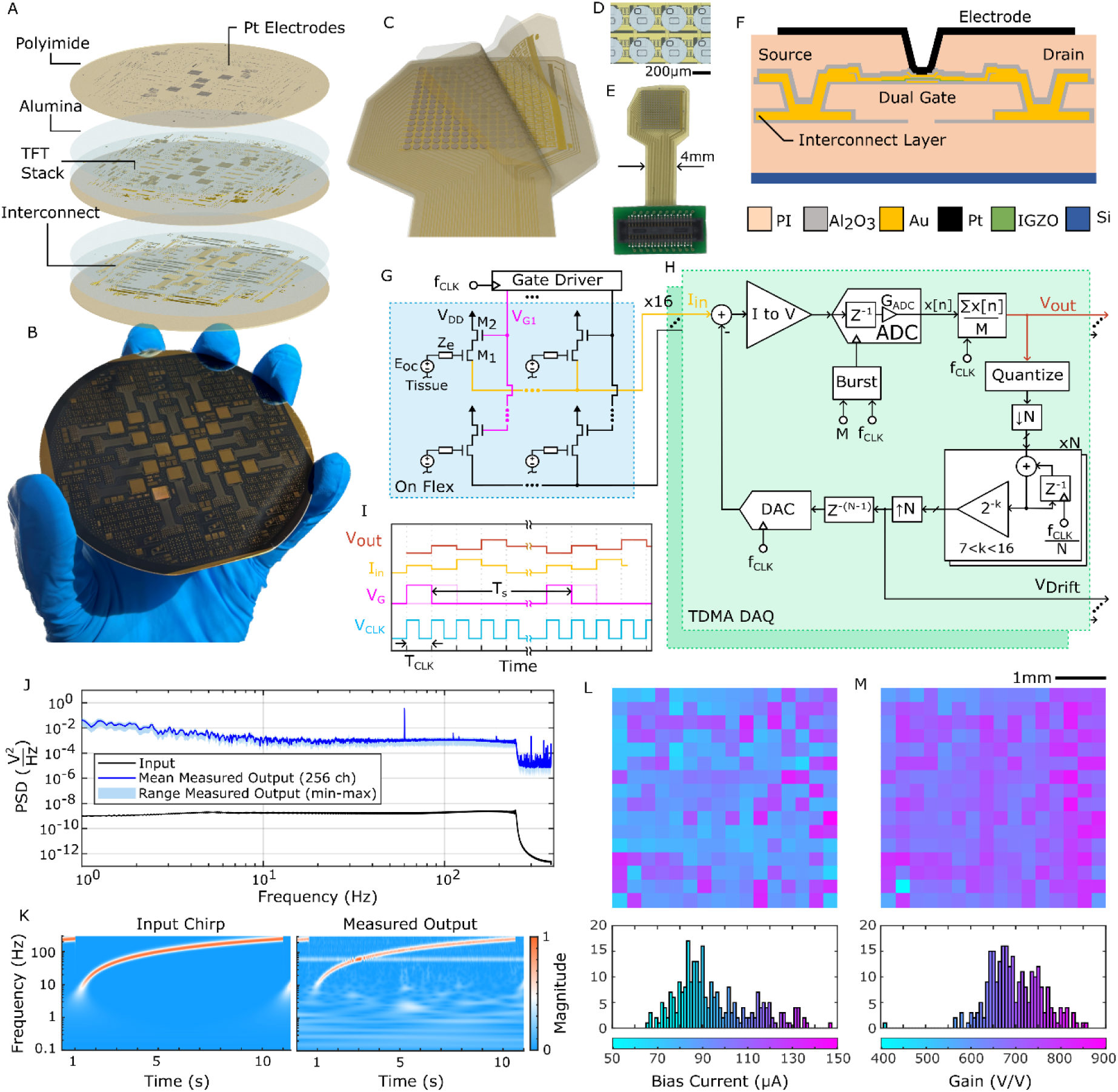
Summary of proposed active neural interface technology stack based on a-IGZO TFTs. A-B) image of 4-inch wafer with exploded view showing layer stack-up containing organic and inorganic encapsulation layers surrounding interconnect metals, TFTs, and top electrode arrays. C) Rendering of 16×16 array of electrodes with pertinent layers separated for easier viewing. D-E) Photographs of 16×16 array bonded to breakout printed circuit board with zoomed-in optical microscope image of the top-view of a few pixel circuits and their corresponding electrodes. F) Cross section rendering representing the key features of the stack-up, including the interconnect layer, the dual-gate TFT, and the sensing electrode. G-I) schematic and block diagrams shown together with simplified timing diagram to demonstrate the proposed transconductor pixel circuit and TDMA shared data acquisition system (DAQ). J-K) power spectral density and corresponding scalograms shown for benchtop measurements in saline taken with an input reference chirp signal. L-M) corresponding measured bias current and calculated gain histograms and heatmaps for all 256 channels of data presented in panels J-K. Measurements demonstrate 100% yield arrays with high process uniformity despite a lack of in-pixel compensation for device characteristics variation.

The fabrication methodology seamlessly integrates TFTs, capacitors, and metal routing with atomic layer deposited (ALD) aluminum oxide thin-film encapsulation layers (Supplementary Fig. 6). Multiple ALD Al_2_O_3_ barriers (≥ 30nm each) conformally coat the active electronics, and polyimide layers serve as the primary structural substrate and biotic interface material. This hybrid approach exploits complementary material properties: polyimide provides mechanical compliance, planarization, and bulk insulation, while the ceramic barriers suppress moisture permeation due to their dense structure and low water vapor transmission rate. Inorganic trenches patterned between polyimide layers enhance inter-layer adhesion and create barriers between active regions and array boarders, mitigating delamination risks and improving mechanical robustness. Arrays were fabricated on 4-inch carrier wafers containing multiple test coupons (Supplementary Fig. 7), then mechanically released in an electrostatically controlled environment and bonded to a breakout board with a minimal-footprint top-side mezzanine connectors (Fig. 1A-E). A representative cross section of a TFT with all encapsulated metal traces and the top-side electrode is shown in Fig. 1F.

The 256-channel arrays implement an active row-column addressing architecture arranged as a 16×16 matrix of platinum electrodes with on-site “pixel” driving circuits beneath each electrode. This on-site active sensing scheme reduces the interconnect bottleneck from O(n) for passive arrays to O(√*n*) - a critical enabler used in the display field to scale modern mobile displays to millions of pixels. Each pixel consists of a sensing TFT (M1) operated in saturation as a transconductor, a switching TFT (M2) for column selection (Fig. 1G), and in later iterations of the technology, a pseudo-resistor and coupling capacitor for bias stability and DC isolation (Supplementary Fig. 8, 9). Column address lines sequentially enabled pixels while parallel row readout lines simultaneously measure current outputs from all 16 rows through high-gain, low-noise transimpedance amplifiers, thereby enabling sequential time division multiple access (TDMA) measurements. A critical innovation that provides an additional avenue for future optimization is the integration of digital feedback: a low-pass digital filter continuously measures and subtracts the DC bias current component at each TIA input from each individual pixel, dramatically increasing dynamic range and enabling per-pixel calibration of operating points. Optimized custom firmware and software handled data acquisition, packaging, transmitting, and logging of the output and drift data shown in Fig. 1H for all 16 parallel channels. This mixed-signal approach significantly reduces power consumption while maintaining high transconductance (Fig. 1H-I). Combined with burst sampling at elevated sampling rates during each column period to reduce aliasing noise, it achieves ∼1ksps per channel with measured per-pixel end-to-end gains exceeding 800 V/V for nominal pixel operating currents of ∼90uA (Fig. 1L-M). It provides a stable mechanism for tracking DC and low-frequency signals associated with TFT operation and drift together with the inherent open-circuit potentials of each electrode (see V_Drift_ in Fig. 1H). This general approach has been pursued in recent works and, when combined with the present mixed-signal architecture, has proven to be a promising avenue for resolving the noise-aliasing problem as well as lowering power scaling with higher channel counts^25^.

Benchtop characterization in phosphate-buffered saline using broadband chirp excitation confirmed a uniform frequency response from DC to 250Hz with minimal channel-to-channel variation (Fig. 1J-K). Spatial mapping of time-averaged pixel currents revealed a relatively large stochastic spread of bias currents (the range varied based on bias voltages), pointing to the need for future pixel architectures to address compensation of device process variation (i.e. threshold voltage variation) rather than relying on downstream digital normalization techniques such as Z-scoring. The system demonstrated accurate signal reconstruction through time-division multiplexing. For example, the power spectral density calculations shown in Fig. 1J-K indicate a measured total input-referred RMS noise of ∼60µVrms. These benchtop validations established the electrical functionality and uniformity of the platform prior to *in vivo* recordings.

### Thin Film Encapsulation and TFT Optimizations for Long-term Active Implants

Hermetic encapsulation represents one of the most significant challenges for flexible active implants, as conventional thin-film polymers exhibit water vapor transmission rates (WVTR) orders of magnitude higher than their bulk-material counterparts. We developed a thin film encapsulation strategy employing polymer substrates and encapsulation layers together with conformal atomic layer deposited high-quality ceramic inter-layers which encapsulate sensitive electronic devices within the FIC. To validate long-term reliability of our hybrid ALD Al_2_O_3_/polyimide encapsulation strategy, we conducted accelerated aging studies at 50°C in phosphate-buffered saline solution with continuous +5V DC bias stress applied between the metal patterns and the surrounding saline over a 388-day period (Fig. 2 A-H). Three encapsulated metal test patterns (Fig. 2A) were monitored via electrochemical impedance spectroscopy throughout the accelerated aging study (Fig. 2G). As shown in Fig. 2H, the impedance magnitude of the insulation at 1kHz declined gradually over the year-long study. Based on fitting a linear degradation model to this data and extrapolating to a failure threshold of one order of magnitude below the initial 1kHz impedance magnitude, we project a lifetime of ∼15.4 years at the 50°C and +5V stress conditions. Applying an Arrhenius acceleration factor of ∼2.5x yields an estimated operational lifetime exceeding ∼38.5 years - far surpassing the requirements for clinical semi-chronic neural interfaces and demonstrating a clear translational pathway for the proposed thin film encapsulation methodology.

**Figure 2.**
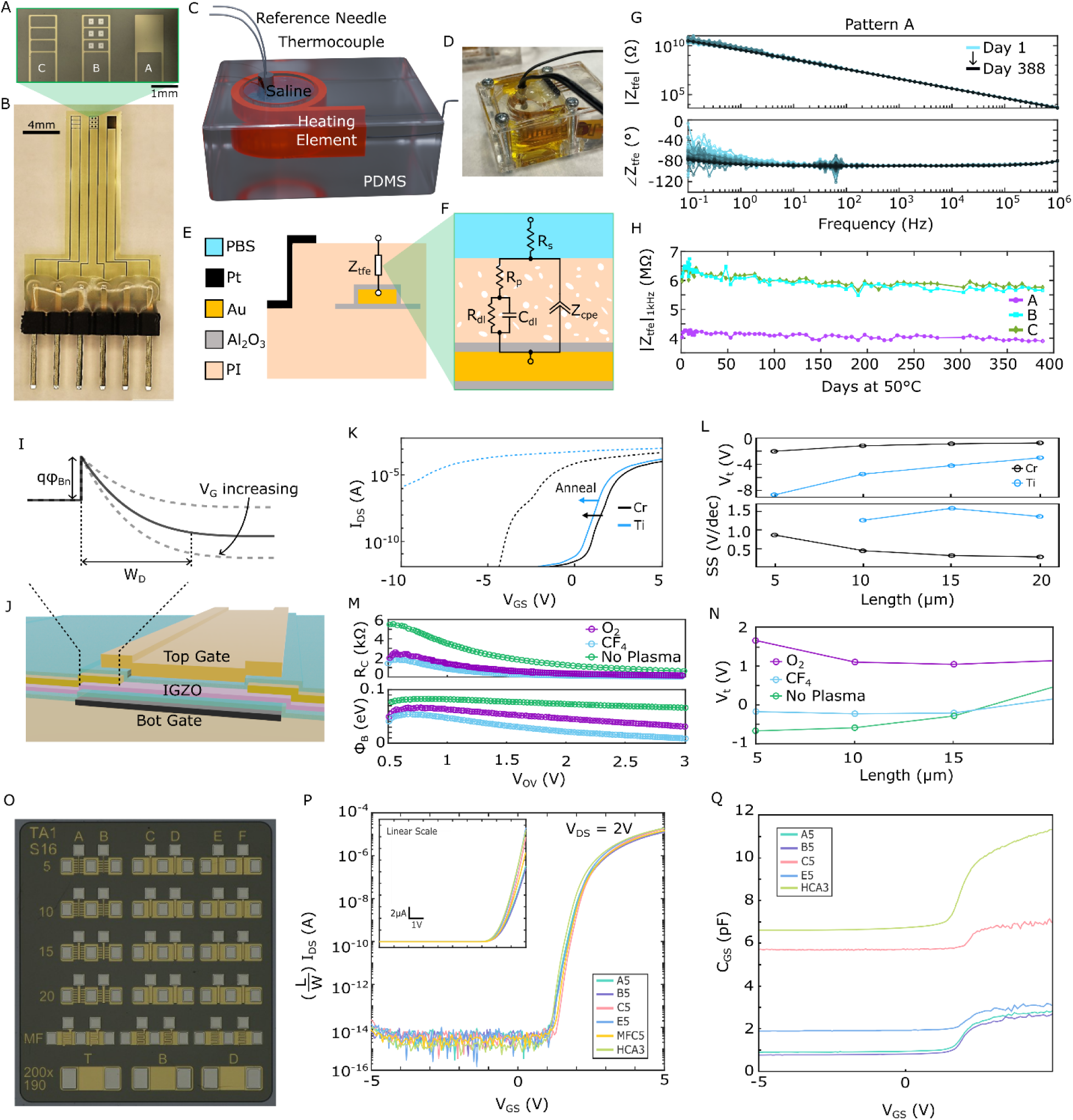
Overview of FIC technology developments. A-B) photograph and optical microscope image inset for encapsulation test samples used in accelerated aging benchtop study. C-D) rendering and corresponding photograph of a PDMS-based accelerated aging soak chamber consisting of a heated saline well wherein the sample shown in panel B was immersed for over one year at 50°C. E-F) Corresponding cross-sectional rendering showing the expected insulation impedance of the imbedded metal leads, together with dummy electrode contacts which were used to assess morphological impacts of thin film encapsulation failure. G-H) Electrochemical impedance spectra and corresponding 1khz impedance magnitude plotted over the duration of the 388-day study. This study was conducted with a 5V DC bias applied across the insulation shown in panels E-F to simulate worst-case scenario of constant operation. I-J) TFT source/drain contact location and corresponding energy diagram as a function of gate bias. K) Transfer curves measured for samples with Cr and Ti contacts before and after annealing. L) Corresponding device characteristics as function of channel length after annealing steps. M-N) Details on gate bias dependent contact resistance and barrier height highlighted for three plasma treatment conditions along with corresponding threshold voltages. O) Microscope image of test coupons used for reported measurements. P-Q) Transfer curves and capacitance-voltage curves for a set of different TFT architectures with details reported in Supplementary Fig. 14-16. Legend in panel P shows code corresponding to test coupons like that shown in panel O, where the letter corresponds to the column code and the number corresponds to the channel length.

Scanning electron microscopy of the age-accelerated samples revealed surface contamination from the phosphate buffer and degraded thermocouple (Supplementary Fig. 4), but no visible delamination, cracking, pinhole defect propagation, or metal corrosion of the encapsulated traces was observed. The ALD alumina layers remained intact with no morphological changes evident, and the platinum dummy contacts from Pattern B (shown in Fig. 2A) appeared in good condition, with only occasional slight peeling at the edges of the contacts which was likely a result of surface scratching caused by intermittent handling during the study. The demonstrated longevity establishes that thin-film encapsulation, when properly engineered with high-quality conformal multi-layer ceramic/polymer stacks, can provide hermeticity at a level of robustness appropriate for implantable electronics applications while maintaining the mechanical compliance essential for highly scalable neural interfaces.

Notably, achieving this robust encapsulation necessitated process temperatures up to ∼200°C during the final alumina passivation and polyimide curing steps (included thin film encapsulation material and processing characteristics are summarized in Supplementary Table 1). This thermal budget initially caused significant instabilities in the TFT characteristics, motivating comprehensive device-level optimizations. We discovered that titanium source/drain contacts, while often used in industry because it provides a suitable work function alignment to IGZO, acted as oxygen and hydrogen getters during thermal processing, notably extracting oxygen from the IGZO channel and creating threshold voltage shifts by increasing oxygen vacancy density (Supplementary Fig. 10, 11). The estimated Gibbs free energy of formation (Supplementary Table 2) for TiO_2_ (-445 kJ/mol O) compared to that of IGZO (-306 kJ/mol O) thermodynamically favored this oxygen redistribution, resulting in unacceptable threshold voltage shifts during thermal processing. Substituting chromium contacts, which are expected to exhibit lower oxygen diffusivity and a smaller energetic driving force for oxidation (-351 kJ/mol O) dramatically improved transfer curve stability when compared against on the same wafer undergoing identical processing (Fig. 2 K). The chromium contacts also exhibited substantially reduced channel-length dependence on threshold voltage shift compared to titanium (Fig. 2L), which we attribute to a lower lateral diffusion of oxygen vacancies into the channel from the contacts during thermal annealing steps.

We also tested various plasma treatments to the IGZO contacts prior to source and drain metal deposition (Fig. 2M-N & Supplementary Fig. 12-14) and found that the fluorine plasma treatment chemistry resulted in the best stability, lowest gate-bias dependent contact resistance and estimated contact barrier height, and threshold voltages closest to zero. We attribute these desirable traits to the fluorine atoms substituting for oxygen vacancies and forming more stable bonds compared to mobile ions such as hydrogen while still providing a free electron donor to the IGZO film.

Beyond fundamental device fabrication optimizations, we also explored several layouts and device architectures to balance current drive, parasitic capacitance, and routing density, some of which are shown in Fig. 2O (additional details in Supplementary Fig. 15-18). The multi-tap contacts used in Columns A and B of the test coupons shown in Fig. 2O aimed to reduce overlap area by 50% while maintaining mostly uniform fields due to contact current crowding and resulting in >70% of the effective channel width, demonstrating a pseudo-self-aligned TFT architecture without prohibitively high source and drain resistances. The benefit of this approach can be seen in Fig. 2Q, where devices A5 and B5 show substantially lower capacitance, attributed to the lower overlap capacitance. Honeycomb and multi-finger layouts (denoted HCA3 and MFC5 respectively in Fig. 2P,Q) achieved the highest packing density with subthreshold slopes as low as ∼110 mV/dec. Supplementary Fig. 18 shows device architecture breakdown characteristics well-exceeding the targeted maximum bias voltages of ±5V employed in our circuits. These device-level safety margins, together with current-limiting resistors and a ground current detection and auto-shut-off mechanism provide robust operation against the electrical hazards present in chronic implant scenarios. The successful co-optimization of insulation longevity and device stability represents a critical milestone for translating flexible TFT technology to implantable neural interfaces, where the slightest system failure is intolerable.

Finally, building off previous modeling efforts in our lab^25^, we developed a simulation platform which assisted us in determining key biasing and circuit design parameters. Supplementary Fig. 19-23 (along with Supplementary Table 3) showcase this platform along with key calculations that help us to assess the impact of thin film stack-ups, device sizing and biasing, the duty cycling introduced from our TDMA approach, and expected electrode and device noise. These simulations, together with benchtop characterization (additional details shared in Supplementary Fig. 24-27) helped us assess the viability of the approach outlined in Fig. 1 and 2 prior to translating our efforts to acute experiments.

### Acute In Vivo Recording

We validated the active arrays through acute cortical recordings in anesthetized Sprague-Dawley rats during sensory-evoked whisker stimulation experiments (Supplementary Fig. 28). In each case, the array was positioned epidurally over the S1 whisker barrel cortex (Fig. 3A), and contralateral whisker deflections were delivered via air puffs while the array recorded broadband neural activity at ∼1ksps. A critical safety consideration for active implants is thermal burden to tissue. We employed infrared thermography to measure cortical surface temperatures before, during, and after the arrays were set to their maximum pixel current and gain settings (∼140 µS TFT transconductance, or ∼750 V/V system gain). The resulting maximum temperature measurement over the cortical surface was captured for several minutes and subsequently segmented into “Baseline” and “Recording” windows (Fig. 3B-C). The data show negligible thermal rise correlated with the operation of the array, clearly below the 2°C limit for chronic implants. The electrode pixel area (300µm x 300µm) in our array design was 180 times larger than the sensing TFT channel area (100µm x 5µm). The sparsity of heat-generating elements together with the 1:16 time division multiplexing of pixel columns (6.25% duty cycle), limited the resulting average power density at each pixel site, allowing for heat to dissipate over a larger area, thus lowering the tissue thermal rise. Our electrothermal simulation results also indicate that the lateral heat spreading through the alumina encapsulation layers likely helped maintain thermal loads well within safety margins.

**Figure 3.**
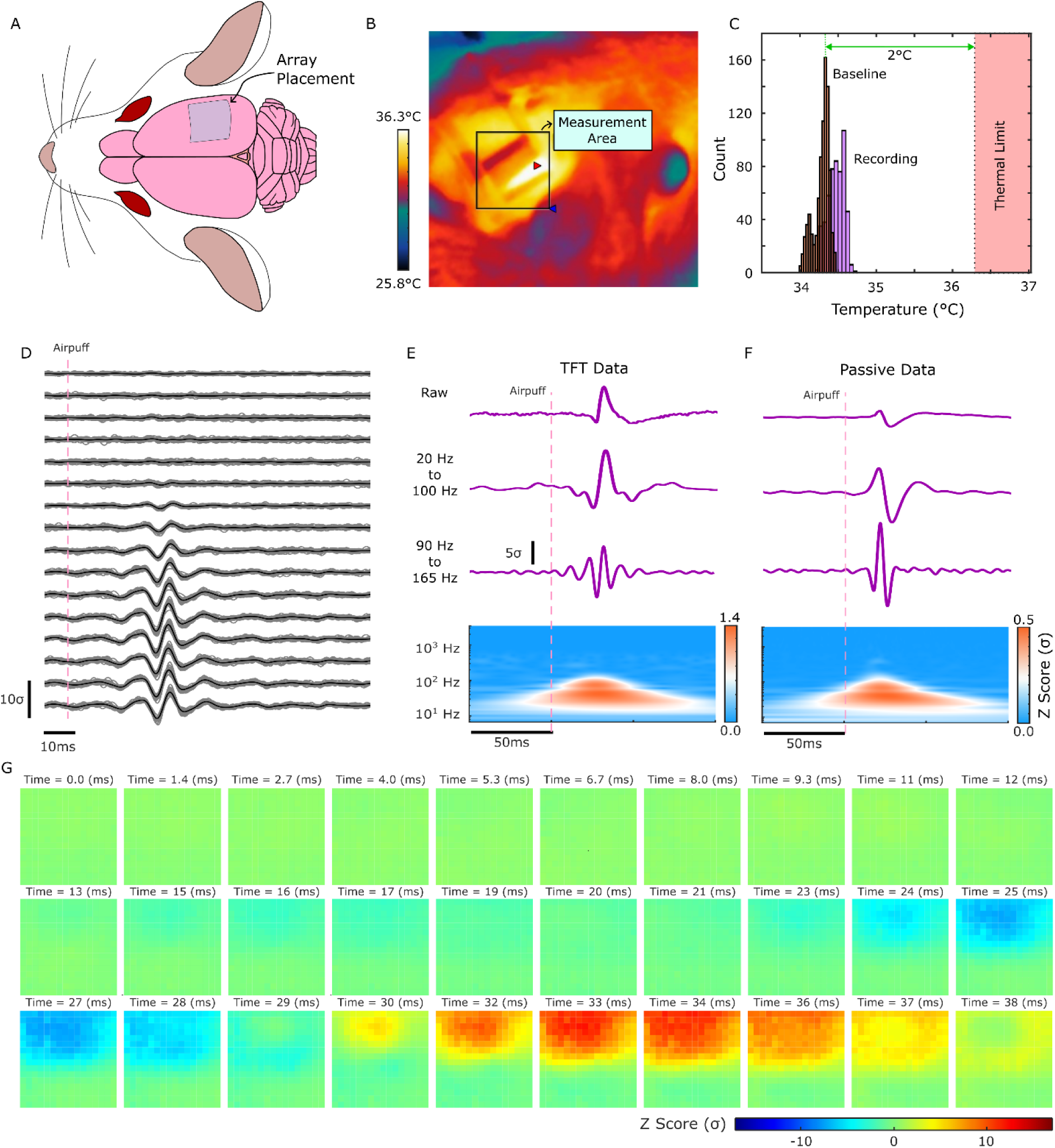
Acute recording of sensory-evoked responses from whisker stimulation. A) Diagram of array placement on rat sensory barrel field. B) infrared surface thermography showing measurement area corresponding to panel A location. C) Corresponding histogram of maximum surface temperature for baseline versus recording during a measurement with maximum bias current settings to determine worst-case thermal burden on cortical tissue. D) Waterfall plot from the middle column in the array showing the z-scored band-pass filtered trials in gray with trial-averages shown in black. E-F) Comparison of an electrode recording with raw and filtered data for the TFT-based active sensing array and an identical passive array. Similar digital filter designs were used to process both datasets to provide a fair comparison, and z-scores were calculated after filtering so as to provide bandwidth-dependent z-scoring details. Corresponding scalograms of the “raw” data are shown below the time-domain traces. G) tile plot heatmaps of the TFT array’s band-pass filtered trial average evoked responses.

After establishing safe thermal loads under maximum gain and operating power conditions, we captured sensory evoked potentials from another array and compared the results to a comparable passive array with equivalent planar platinum electrode size and pitch. The trial averaged evoked responses are shown in Fig. 3D-G. A waterfall plot of band-pass filtered trials and their resulting trial averages are shown in gray and black traces respectively for the middle of the array in Fig. 3D. The corresponding trial averaged z-scored waveforms for a single responsive channel from both the FIC-based array and the passive comparison array (which connected to an Intan Technologies recording system) is plotted in Fig. 3E-F. Both datasets were captured from the same animal with identical electrode contact geometries and similar cortical coverage. The TFT array waveforms (Fig. 3E) are inverted and have slightly different shapes relative to the corresponding passive array waveforms (Fig. 3F). We attribute these differences mostly to the inverting nature of the transimpedance amplifier and the system transfer function for the active neural interface system. While the differences could be compensated via postprocessing that utilize estimations of each per-pixel system transfer function, we decided that without more rigorous system calibration in place to determine the precise deconvolution across frequencies, a z-scoring methodology would provide a fairer method of comparison. In the frequency domain, it becomes apparent that the captured evoked cortical potentials for both electrode arrays share a similar bandwidth up to ∼200Hz, with the majority of the gamma and higher-frequency signal content beginning roughly 20ms after stimulus and ending roughly 40ms after stimulus. Notably, the passive system appeared to have worse z-scoring at lower frequency which we attribute to both the difference in signal conditioning mechanisms and the difference in routing and loading (noting that the high input impedance of the sensing TFT gate minimizes signal attenuation and provides on-site buffering). The passive system also appeared to have improved signal to noise ratios (as evident by increased z-scores in Fig. 3F) within the high-gamma band. This is likely a result of noise-folding from aliasing due to the on-site multiplexing together with the lower sampling rates employed in our custom acquisition system as compared to the passive recording system which used the Intan Recording Controller, sampling at 20ksps. Despite these limitations, we see robust sensory responses for our active neural sensing arrays with z-score peak responses exceeding 8σ in maximally responsive channels and occurring within the expected 20-30ms post-stimulus delay time.

The spatiotemporal response propagation presented in the tile plot in Fig. 3G also validates the functionality of our active sensing scheme. We see clear spatiotemporal propagation of the whisker barrel field response to stimulation, with negligible observed spatial artifacts. These results establish that the TFT-based platform can capture high-fidelity neural signals with spatiotemporal resolution comparable to state-of-the-art passive arrays while offering a viable pathway to dramatically increased channel counts through the reduced interconnect requirements.

### Chronic Recording in Sprague-Dawley Rats

To assess translational viability, we conducted a month-long chronic implant study in Sprague-Dawley rats with the active array positioned epidurally over the S1 barrel cortex. The chronic implant housing shown in Fig. 4A was designed to house the array. The implant consisted of the same 256-channel design used in the acute study pre-assembled into the housing and implanted using a combination of medical-grade Metabond bone cement, Kwik-Cast silicone sealant, and Gelfoam (Supplementary Fig. 29-30). A critical design feature enabling chronic implant success was the implementation of pseudo-resistor-based bias stabilization and ESD protection. Between recording sessions, all array terminals were electrically shorted via a removable connector shunt, preventing static charge accumulation across the gate dielectrics - a failure mode we identified during earlier pilot studies when low-humidity vivarium conditions generated substantial electrostatic fields within the insulating cages.

**Figure 4.**
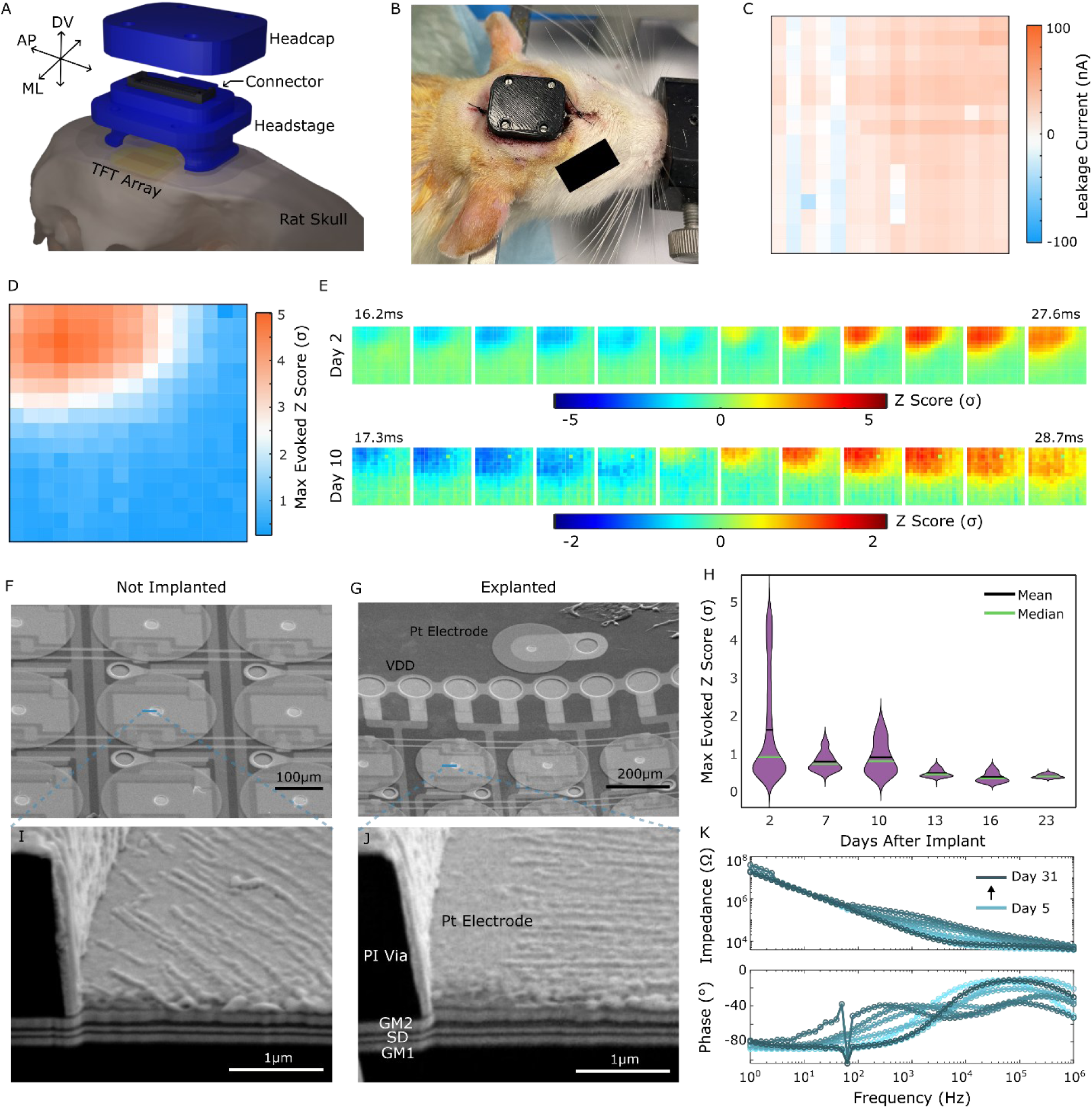
Overview of chronic implant study showcasing long-term viability of our active neural interface. A) 3D rendering of the chronic implant. B) image of the rat post-implantation showing the headstage and headcap. C) Exemplary leakage current measurement taken prior to each recording, plotted spatially for all pixels. D) maximum evoked z-score within response window for trial-averaged high-gamma activity on day 2 of study. E) Days 2 and 10 tile plots for evoked responses showing clear evoked responses corresponding to whisker barrel activity. F-G) Scanning electron microscopy of post-mortem explanted array compared against a non-implanted array (fabricated on same sample wafer but not implanted) with corresponding cross-sectional FIB imaging demonstrating identical layer stack-up of electrode via sidewall profile and underlying TFT stack. H) Violin plots across implant days showing max evoked z-scores across all sensory-evoked recordings. K) Electrochemical impedance spectra of passive electrode (shown in panel G) throughout duration of implant showing increasing 1khz impedance magnitude resulting from the development of encapsulation tissue between the cortical surface and the electrode array.

Electrochemical impedance spectroscopy of a passive reference electrode (SEM image seen in Fig. 4G) on the array are reported in Fig. 4K and Supplementary Fig. 31 to show a gradual impedance increase over the month-long study (from ∼30 kΩ to ∼60 kΩ at 1kHz), consistent with expected fibrotic encapsulation of the epidural implant. Critically, the insulation impedance of a dedicated encapsulated test pattern (a similar pattern to those shown in Fig. 2A-B) remained capacitive, as predicted by our accelerated aging studies. Furthermore, the pixel performance was assessed through calculated kernel densities of transconductance efficiency across all recordings of the chronic study (Supplementary Fig. 32, 33), indicating similar sensing TFT operating regimes were achievable throughout the experiment. We attribute this in part to the stability of our TFTs owing to the optimizations in fabrication processing conditions discussed above, along with our thin film encapsulation strategy which incorporated high-density ALD-deposited alumina that appeared to effectively protect the oxide TFTs from environmental aggressors.

Array calibration measurements conducted during each recording session demonstrated stable TFT operation, as can be seen in Fig. 4C, where per-pixel leakage current and other relevant characteristics were assessed prior to each recording through a bias sweeping procedure discussed in Supplementary Fig. 25. Sensory evoked recordings performed on days 2, 7, 10, 13, 16, and 23 consistently captured whisker deflection responses with clear spatiotemporal structure (Fig. 4 D-E), though the final recording on day 31 was unsuccessful due to a partially damaged headstage. Trial-averaged high-gamma signals revealed peak z-scores around 5σ in the first recording on Day 2 with some variation and gradual decline across the study duration attributed to the variable nature of evoked sensory responses under anesthesia together with the observed development of fibrotic encapsulating tissue. Nevertheless, the spatiotemporal wave dynamics of evoked responses remained consistent across recording days, with weighted centroids clustering within a ∼400 μm region of the array (Supplementary Fig. 34), with subtle differences in propagation direction and spatial spreading which is believed to be related to variation in whisker stimulation, anesthesia, and encapsulation tissue development between the implant and the cortex. These results demonstrate that the addressing architecture maintained channel independence and spatial fidelity throughout the implant duration, regardless of a weakening signal source due to the natural angiogenesis associated with wound healing.

Post-explant analysis revealed 1-2 mm thick fibrous encapsulation surrounding the cranial window and underlying the array - an expected foreign body response for epidural implants. Despite this tissue growth, the flexible array remained mechanically intact with no visible delamination, cracking, or material degradation. Scanning electron microscopy of the explanted array after thorough cleaning showed surface morphology and cross-sectional layer thicknesses identical to a pre-implant reference sample, with no evidence of Al_2_O_3_ dissolution, polyimide swelling or degradation, or platinum electrode corrosion or delamination (Fig. 4 F-G). This structural preservation after 30 days of *in vivo* exposure, combined with the maintained electrical functionality, strongly suggests that the encapsulation strategy is viable for long-term implantation.

The chronic study represents the first published demonstration of a fully integrated flexible TFT-based active neural interface maintaining functionality over multiple weeks *in vivo*. While the one-month duration is modest compared to multi-year clinical requirements, the combination of projected encapsulation lifetime (>38 years), stable device characteristics, and successful signal capture establishes a foundation for translational development. Key challenges for future work include reducing foreign body response through subdural placement, improving per-pixel gain uniformity through compensation circuitry, and extending the channel count through the combined use of custom driver integrated circuitry and TFT-based shift-register gate addressing.

Towards these goals, we also developed and validated TFT-based dynamic shift-register architectures shown in Supplementary Fig. 35 - 39 which paves the way for future flexibility scalability improvements. In modern-day AMOLED displays, gate-on-array (GOA) approaches have unlocked unprecedented scalability in pixel densities and narrow display panel boarders. The NMOS-only dynamic shift-register architecture provides a feasible pathway for implementing next-generation flexible active neural interface circuits which can scale towards pixel densities comparable to modern-day mobile displays (e.g. millions of pixels).

## Conclusion

We have demonstrated a monolithic FIC platform for active neural sensing that addresses the fundamental interconnect bottleneck that has limited passive electrode arrays. By adapting high-definition AMOLED display technology to neural interfaces, our a-IGZO TFT-based system achieved O(√*n*) interconnect scaling (with next-generation scaling methodologies presented in Supplementary Fig. 35 - 39) through in-pixel power amplification and row-column multiplexing while maintaining mechanical compliance with neural tissue. Critical innovations in thin film device fabrication and architectures helped to enable this technology, including contact engineering, surface treatment, and careful attention to layer stack-up design. Year-long accelerated aging studies projected >38-year operational lifetimes for our hybrid ALD Al_2_O_3_/polyimide encapsulation stack, bringing into question the need for bulky metallic or ceramic enclosures used in conventional implantable electronics. Acute and chronic *in vivo* validation in the rat model confirmed safe operation with negligible thermal burden, signal quality exceeding comparable passive arrays, and maintained functionality over 30 days of implantation. These results, to our knowledge, represent the first demonstration of a fully integrated flexible TFT-based active neural interface maintaining chronic functionality.

Compared to silicon-based active neural interfaces, our approach offers distinct advantages in mechanical compliance and substrate-agnostic fabrication, offering the potential to reduce implant invasiveness over thicker, more rigid alternatives. The monolithic fabrication approach presented here provides an easier path to high-yield commercial fabrication at volume while maintaining full design flexibility for future design optimizations as compared to more heterogeneous fabrication approaches which may require more advanced integration strategies that may compromise hermeticity of encapsulation and set limits on implant size. These advantages make our approach amenable to more traditional panel manufacturing flows which further elevates the likelihood of future clinical translation.

Looking forward, several exciting pathways exist to advance this platform toward clinical viability. First, incorporating in-pixel compensation circuits - such as threshold voltage and bias stress compensation - would dramatically improve gain uniformity and device stability across larger arrays. Second, integrating dynamic shift registers for gate driver addressing, as employed in modern AMOLED displays, could reduce interconnect scaling even further, unlocking the potential for broader cortical coverage with higher channel counts. A third future direction is to directly integrate an ASIC similar to the one our group developed at UCSD^37^ into the flexible substrate through flip-chip bonding to consolidate all analog front-end, digitization, and clock generation into a single ultra-low-profile implantable package. This approach would reduce the entire system size to a highly scalable and easily implantable form factor, though it should be noted that the integration of rigid ASICs directly into the flexible substrate would require significant technological development and introduces many challenges that were discussed in the above introduction section. Finally, if thin film transistor technology ever advances to a point where the mixed-signal front-end circuitry can be reasonably implemented on-flex, this would unlock the potential for fully monolithic implants. Unfortunately, for this ultimate dream to be realized, several technology blocks are still needed, including a thin-film battery or super capacitor and a more advanced complementary thin film transistor technology

## Methods

### Fabrication of FIC

All fabrication was carried out at the UC San Diego Nanofabrication Facility (Nano3) on 4-inch silicon carrier wafers through a three-phase process (Supplementary Fig. 1). Phase 1 establishes the bottom metallization and hermetic vertical interconnects (vias). A 10 µm polyimide film (HD MicroSystems PI2611) was spun-cast onto cleaned silicon wafers, followed by atomic layer deposition of >30 nm Al_2_O_3_ at 200°C using trimethylaluminum and H2O precursors. Chrome/gold bilayers (20/200 nm) were sputtered and patterned via a two-step photolithography and wet etching process. A second Al_2_O_3_ layer conformally passivated the metallization, and a 4.5 µm polyimide layer was spun-cast and cured on top. Vias were dry-etched with oxygen plasma using high pressure (>30 mTorr) to achieve graded sidewall profiles, then passivated with an additional ALD-deposited Al_2_O_3_ layer.

Phase 2 of the fabrication flow formed the dual-gate TFT stack. Chrome/gold (50/100 nm) formed the bottom gate, with gold removed selectively from the channel regions to reduce electric field enhancement on sharp negative line edges. After cleaning with oxygen plasma and heated Remover-PG, a 25 nm Al_2_O_3_ gate dielectric was deposited via thermal ALD with TMA a pre-pulse nucleation recipe used to improve film density and reduce pin-hole formation. Samples were immediately transferred to an ultra-high vacuum RF magnetron sputtering system (base pressure <4e-8 Torr) and slowly brought to a stable processing temperature of 200°C. A 30 nm amorphous IGZO film was deposited at 2 mTorr with an O_2_:Ar partial pressure ranging from 15% to 30% based on desired device characteristics (with roughly a 4%/V linear relationship observed between partial pressure and final threshold voltage). IGZO and bottom gate dielectric were subsequently patterned via wet etching. Source and drain contacts (20 nm Cr or Ti / 50nm Au) were deposited via electron-beam evaporation following a low-power plasma treatment and subsequently lifted off under Remover-PG. Finally, the top gate dielectric (25 nm Al_2_O_3_) and Cr/Au top gate were sequentially deposited and patterned using ALD and sputtering respectively.

Phase 3 completed the encapsulation of the FICs. Another (>30nm) Al_2_O_3_ layer passivated the TFT stack, and after patterning to open up contacts and expose underlying organic layers (to improve adhesion) a final overmold of polyimide was spun-cast and cured under vacuum-oven at 195°C, with carefully controlled ramping and environmental monitoring in place to ensure consistency across fabricated batches. Top vias were dry-etched, and chrome/platinum (20/200 nm) electrode metallization was sputtered and patterned via lift-off. Outlines were defined through final dry-etching, and test coupons were then characterized using the B1500A semiconductor device analyzer prior to mechanically peeling samples from the carrier wafer in deionized water in an electrostatically controlled environment under microscope. Samples were then electrically bonded to custom printed circuit boards through silver epoxy selectively applied to bonding pads through a laser-cut stencil mask.

### Animal Experiments and Surgical Procedures

All animal experiments were approved by the UCSD Institutional Animal Care and Use Committee (IACUC), and a summary table of these experiments is included as Supplementary Table 4. Adult male Sprague-Dawley rats were used in both the acute and chronic studies. Craniotomies were performed in a sterile environment to expose the S1 sensory barrel field. General anesthesia (4% isoflurane gas) was administered to the rat through a stereotaxic nose-cone prior to surgical preparation, and titrated down to 1.5-2.5% throughout the craniotomy (adjusting as needed based on heart rate and respiration rate). Local anesthetic (lidocaine) was also administered in the scalp prior to incision. To mitigate anesthetic impact on sensory-evoked activity animals were transferred from isoflurane to a ketamine/xylazine mixture after the craniotomy procedure. Body temperature was maintained through a surgical heating pad and monitored throughout the experiment and recording. Recordings were performed in a faraday cage which housed the custom acquisition equipment.

In the chronic study, the sterile 3D-printed enclosure which housed the implanted array was made of ABS plastic and was secured to the rat skull using Metabond dental cement and sealed closed using Gelfoam, Kwikcast silicone, an additional 3D printed cover-plate, and a final Metabond layer. The scalp was sutured closed around the enclosure to reduce the wound area and improve the surgical outcome. A topical antibiotic was applied to the scalp and edges of the enclosure after suturing (and as needed after the surgery) and an anti-inflammatory (Carprofen) was administered to the rat after surgery. Buprenorphine (Ethiqa XR) was also administered during and after surgery. Rats were monitored for behavioral changes and weight loss throughout the study. At the end of the studies, rats were euthanized under deep anesthesia using a 120 mg/kg sodium pentobarbital intraperitoneal injection.

### Data Acquisition and Analysis

A fully custom battery-powered acquisition system was developed to record from the TFT-based active neural interfaces. The system consists of a dedicated microcontroller running optimized firmware that controls voltage regulators, parallel ADCs and DACs, gate driver circuits, and an electrically isolated transistor-transistor logic (TTL) synchronizing output. Arrays were connected via mezzanine connectors and a flat flexible cable interposer to this external driver circuitry which generated sequential column address clocks while simultaneously measuring pixel currents through 16 parallel transimpedance amplifiers (5.1 MΩ gain). Data were streamed in real-time via USB 2.0 to a recording laptop running custom C++ software. A worker thread serviced the data queue to maintain negligible USB latency and ensure zero data loss. Data were logged using memory-mapped binary files, enabling simultaneous acquisition and visualization of both the 256-channels of 16-bit recording data and 256-channels of 16-bit low-frequency drift data, and each parallel sample was packaged with timestamp and frame number header data at the microcontroller-level to ensure consistency in acquisition sampling frequency and to help check against data loss. The graphical user interface displayed real-time waterfall plots and spatial heatmaps of both high-frequency signals and pixel DC bias currents. Auto-calibration sequences, system settings, and hardware-level debugging data were logged alongside neural data and case notes to provide full visibility into system performance and to streamline recording sessions.

For sensory-evoked experiments, the TTL output triggered a benchtop air-puff system (PV830 Pneumatic Pump) to deliver contralateral whisker deflections at 20 psi with 0.5 Hz frequency (2 second inter-trial interval). Trigger timing was automatically logged relative to acquisition timestamps. Each recording file typically consisted of up to 2 minutes of continuous acquisition with an 11 second baseline preceding the first stimulus onset. Acute experiments lasted several hours, and chronic recording sessions were limited to 1.5 hours to minimize anesthetic duration.

Raw data were resampled using spline interpolation (effectively implementing a digital fractional-delay filter) to eliminate phase offsets between time-division-multiplexed columns, producing a uniform time vector at 16 times the per-pixel sampling rate (∼6.4 kHz effective sampling frequency). Digital filtering was implemented in MATLAB using the following cascade: 2nd-order Butterworth bandstop filters at 60 Hz (59-61 Hz), 120 Hz (119-121 Hz), and 180 Hz (179-181 Hz) to remove line noise and harmonics, plus a bandstop at Fs/4 to suppress aliasing artifacts. Broadband gamma (20-100 Hz passband, 10 Hz and 110 Hz stopbands) and high-gamma (90-165 Hz passband, 55 Hz and 200 Hz stopbands) filtering employed Butterworth bandpass designs with 1 dB passband ripple and 10 dB stopband attenuation. All filters were applied using zero-phase digital filtering (filtfilt) to prevent temporal distortions. Data were z-scored on a per-pixel basis using session-wide mean and standard deviation to enable comparison across varying gain settings.

Trial-averaged evoked responses were calculated by segmenting filtered data into stimulus-aligned windows. Spatial heatmaps displayed maximum z-scores during 0-50 ms post-stimulus intervals. Time-frequency analysis employed continuous wavelet transforms to generate scalograms. Weighted centroids of evoked activity were calculated on thresholded binary masks (threshold = maximum pixel value minus 2 standard deviations across pixels).

Electrochemical impedance spectroscopy measurements were acquired using a Gamry Interface 1000E with potentiostatic mode, 10 mV RMS AC excitation, 0V DC relative to open-circuit potential, and frequency sweep from 1 MHz to 1 Hz. Measurements were performed in saline benchtop experiments, at each chronic recording session, and post-explant for passive reference electrodes and encapsulated test patterns. Continuity measurements were also monitored across the encapsulated test patterns to ensure bonding pad viability throughout the chronic study.

## Acknowledgements

This work was performed in part in the San Diego Nanotechnology Infrastructure (SDNI) of UCSD, a member of the National Nanotechnology Coordinated Infrastructure, which is supported by the NSF (grant ECCS1542148). The authors would like to acknowledge support from the Nano3 staff, in particular Patrick Driscoll, who provided countless hours of technical support and insight into the fabrication equipment required for this research. The authors would also like to acknowledge support from Dr Hoi Sang U of UC San Diego for his guidance in the animal studies and superb surgical training. Finally, the authors would like to acknowledge the pivotal impact of Dr. Hongseok Oh, currently at Soongsil University, who not only provided training and guidance, but also helped lay the groundwork for the fabrication of IGZO thin film transistors implemented in this work. The research presented here built on nearly a decade of know-how that all began with Dr. Oh and his published works on zinc oxide pressure sensors.

## Author contributions

Conceptualization: A.M.B, S.A.D, I.G.

Method Development: A.M.B., R.V., K.J.T., S.A.D., I.G.

Fabrication: A.M.B., R.V.

Data Collection: A.M.B., R.V.

Data Processing: A.M.B

Funding Acquisition: A.M.B., S.A.D., I.G.

Project Administration: S.A.D., I.G.

Manuscript writing: A.M.B., S.A.D.

Manuscript Reviewing & Editing: A.M.B., R.V., K.J.T., S.A.D., I.G.

## Data and Code Availability

All data can be found in the main text or supplementary materials. Other data that support the findings of this study are available from the corresponding author upon reasonable request. The code used for processing the data are also available from the corresponding author upon reasonable request.

