## Supplementary Figures for "A Thin Film Transistor Backplane for Scalable Chronic Neural Interfaces"

### Display-Inspired Flexible Thin-Film Electronics Enable Scalable, Chronic Active Neural Interfaces

#### Supplementary Figures

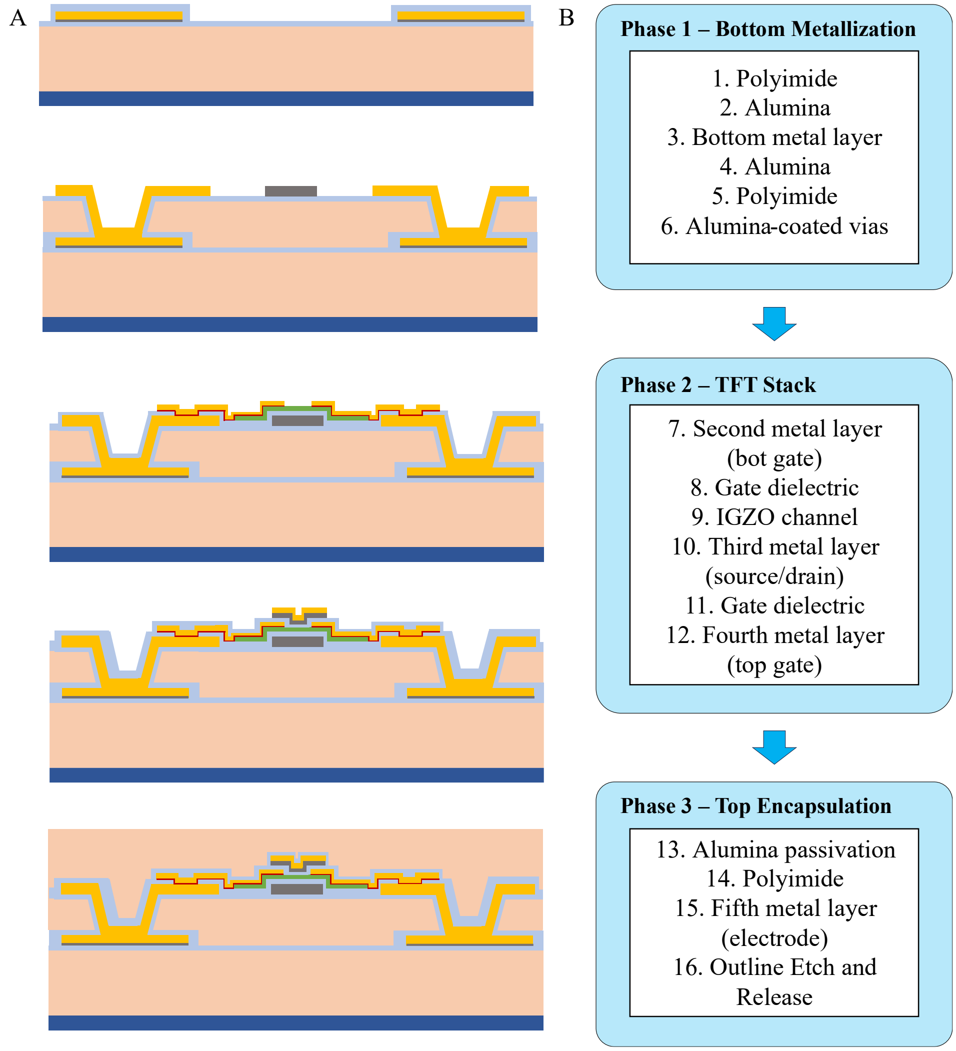


Figure 1 Overview of our fabrication process for a-IGZO based flexible thin film integrated circuits. A) Cross-sectional view of key stages throughout fabrication process. B) Corresponding phases in fabrication listed in chronological order.

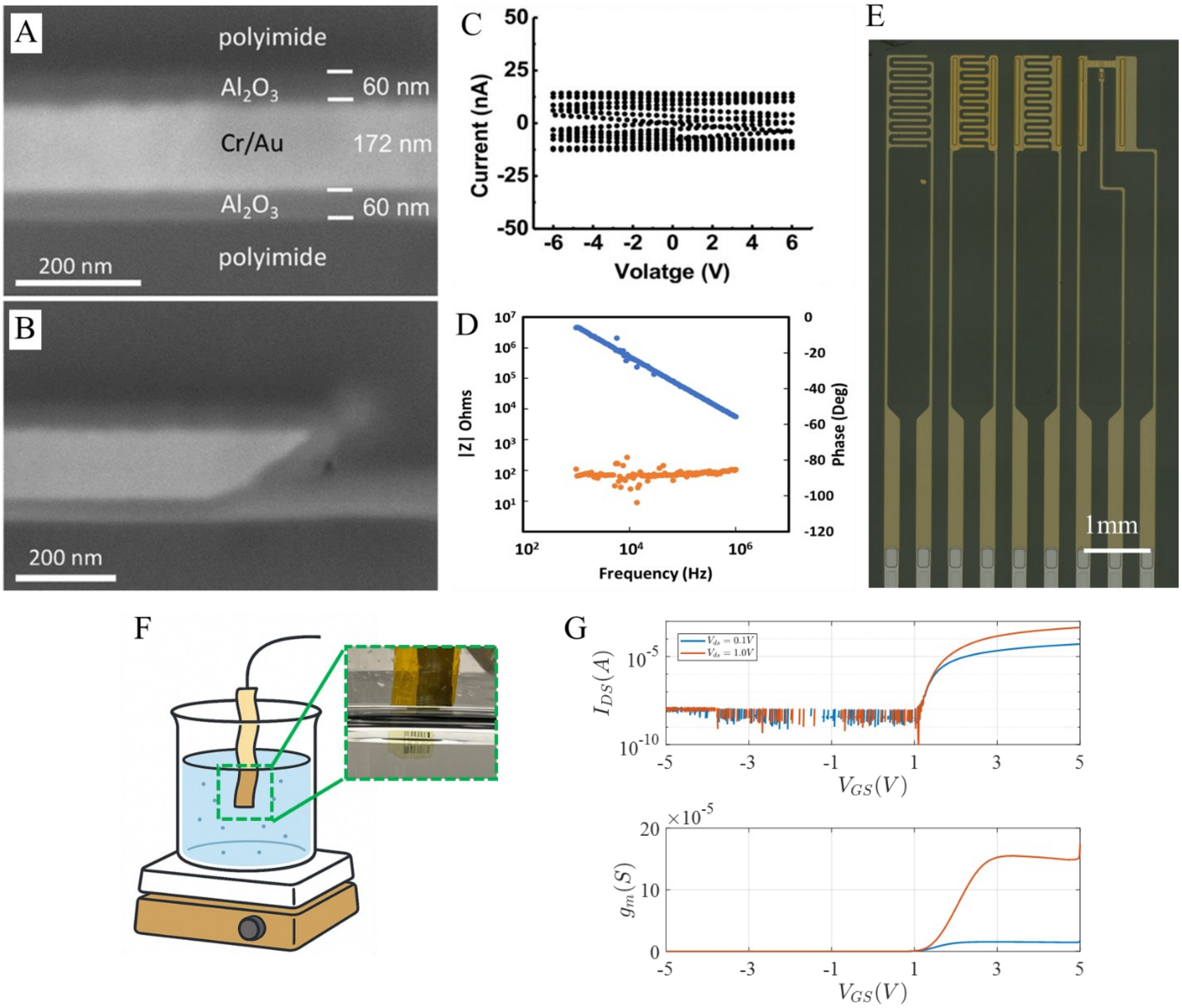


Figure 2 Soak test data for passive interdigitated patterns and a single TFT. A-B) Representative cross sectional imaging showing the highly-conformal encapsulation of metal layers with ALD-deposited alumina. C) Cyclic voltammetry measurements for > 50,000 cycles at +-6V amplitude showing negligible current through insulation (noise floor). D) Potentiostatic electrochemical impedance spectra showing capacitive impedance of insulation. E) Optical imaging of test pattern, including two interdigitated patterns on different metal layers together with a single encapsulated TFT pattern. F) Representation of initial soak test measurement setup together with image of submerged sample, and G) corresponding transfer curves (measured while sample submerged using “fast-mode”, so off current was noise-floor limited) for TFT after 24 hours of soaking in saline.

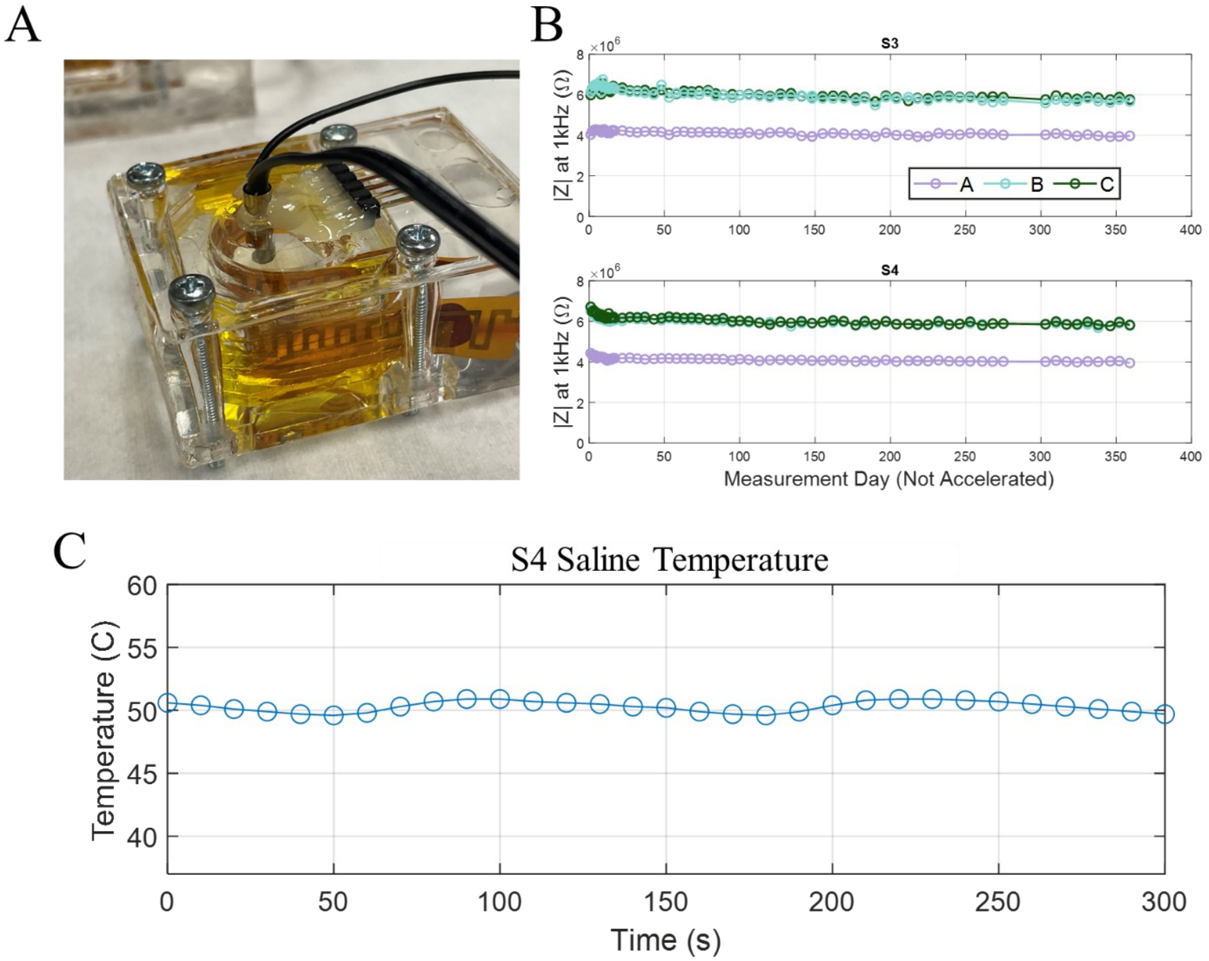


Figure 3 Details relating to accelerated aging effects on 1kHz impedance magnitude for all samples under investigation. A) Photograph of one of the samples when installed in the PDMS saline soak test chamber. B) 1kHz impedance magnitude plotted for all three patterns for samples 3 and 4 plotted over the full year of accelerated aging, where the x-axis was shown in real time, not the estimated accelerated time. C) A representative measurement of saline temperature showcasing a mean temperature of 50°C with minimal thermal ripple introduced from the PID controller

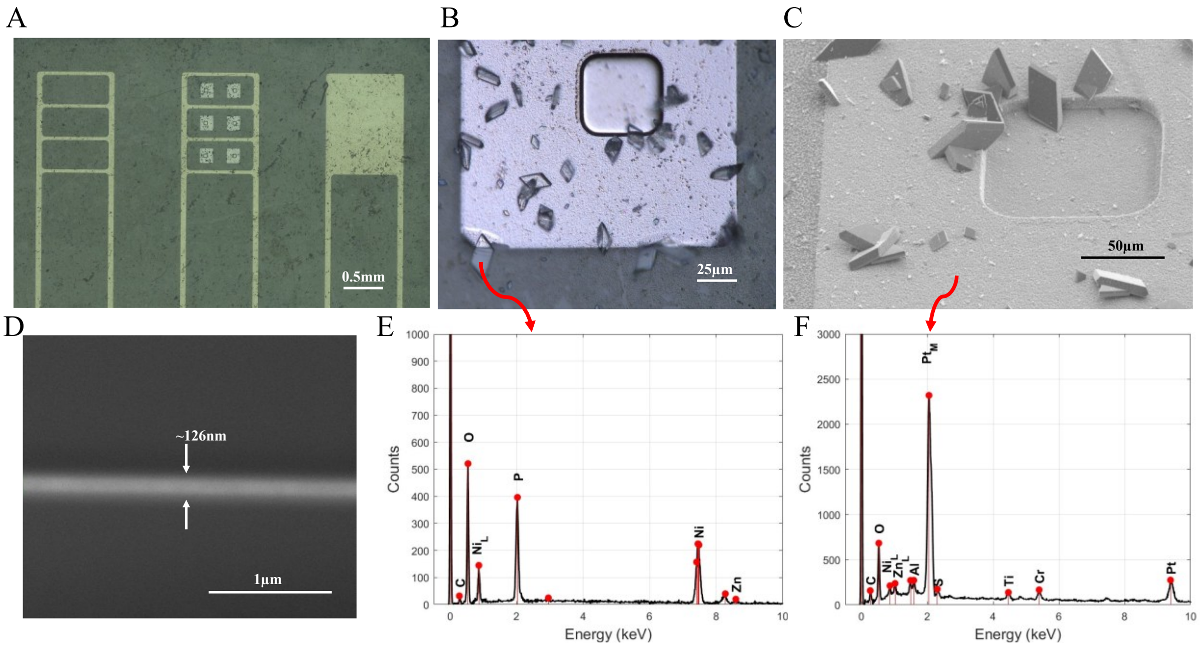


Figure 4 Surface imaging and EDX analysis after year-long aging test for S4 sample. A-B) Optical microscopy imaging showing in-tact patterns and platinum dummy electrode surface in pattern B. Observed crystalline-like residue scattered throughout the surface which was initially believed to be residual salt crystals. Red arrows indicate locations of corresponding EDX measurements shown in panels E-F. D) Cross sectional SEM image of pattern A showing the expected ~120nm thin film of gold remaining well-encapsulated by insulation, with no obvious morphological abnormalities observed. E-F) EDX measurements with overlaid estimates for elemental composition. Results in E indicate the presence of nickel, zinc, and phosphorous and panel F shows platinum surface primarily composed of elemental platinum and chromium (which was used as an adhesion promotor).

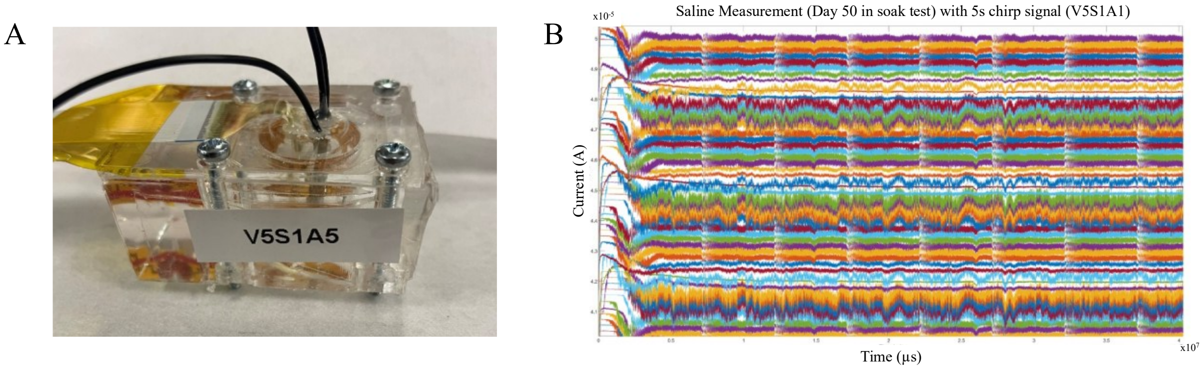


Figure 5 Brief overview of system saline testing demonstrating signal transduction of chirp signals after 50 days accelerated aging.

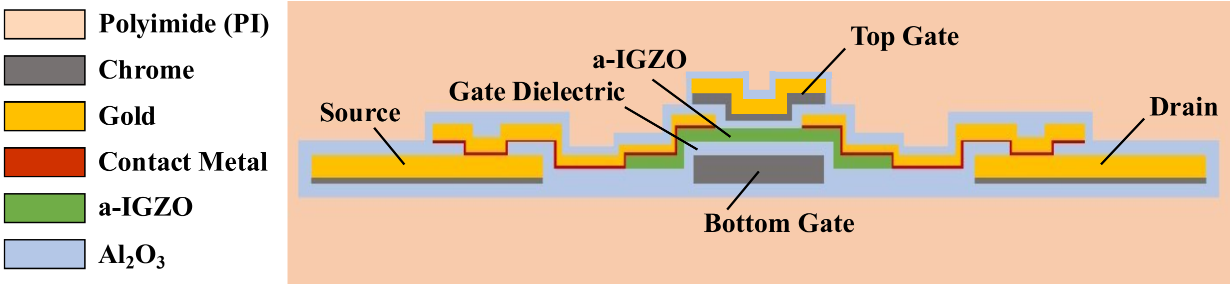


Figure 6 Close-up cross-section of dual-gate a-IGZO TFT with labels for key features.

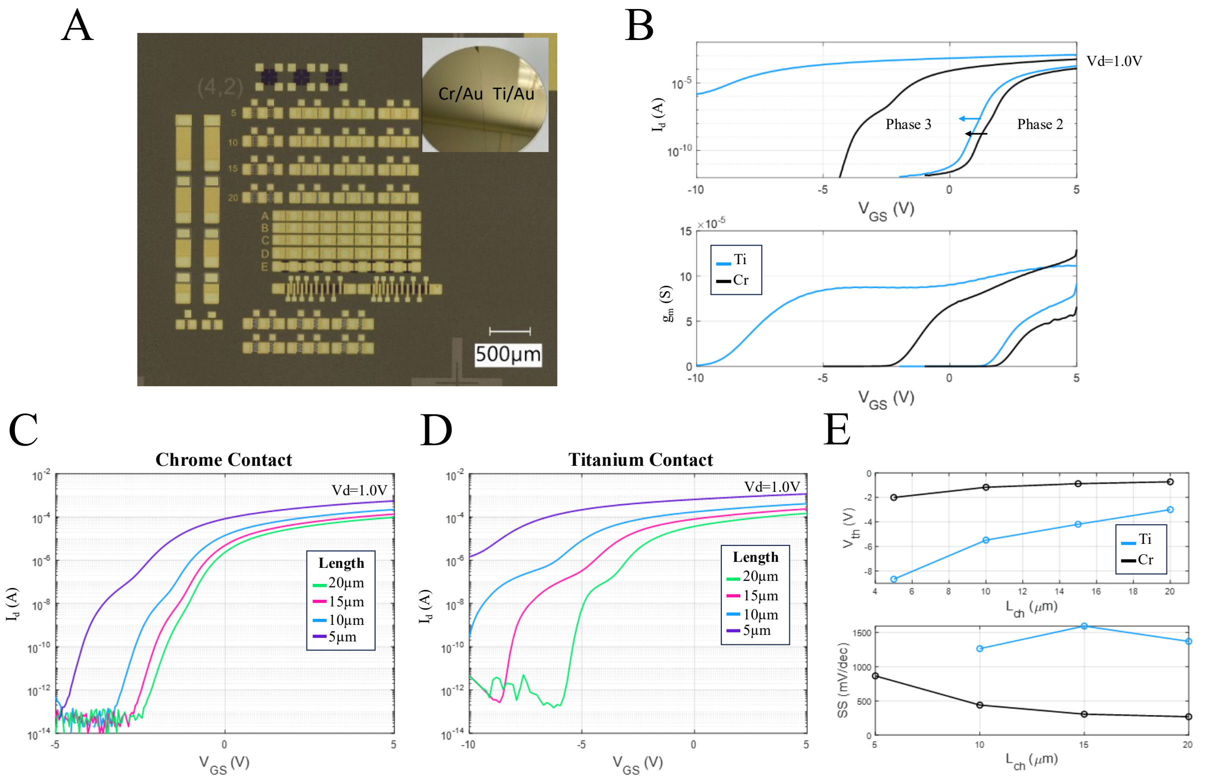


Figure 7 Overview of results from contact metal study. A) Optical die micrograph of a single test coupon with top-right inset showing 4-inch wafer after depositing contact metals through electron beam deposition. B) Transfer curves and corresponding transconductance plotted for representative bottom-gate TFT patterns (W=100μm, L=5μm) for both contact metals across different phases of fabrication. C-D) Transfer curves for varied lengths at the end of phase 3. E) Threshold voltage and subthreshold slope extracted from the transfer curves in C-D. Threshold voltage was extracted from low-Vds transfer curves using an automated regression fitting along with capacitance densities measured from nearby MIM capacitors shown on the left-hand side of the optical die micrograph in panel A.

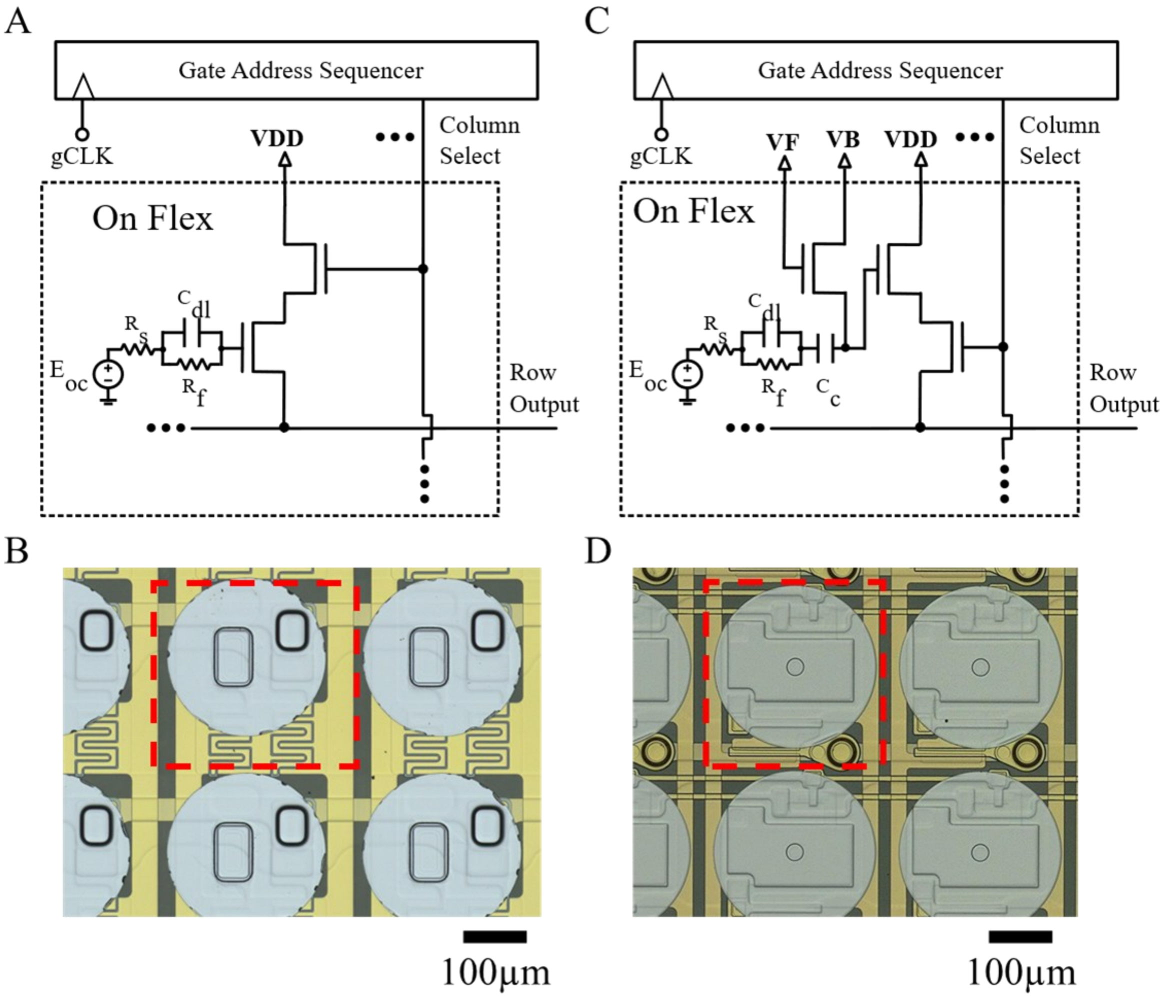


Figure 8 In-pixel time division multiplexing using IGZO TFTs. A-B) Pixel circuit and corresponding optical image for depletion-mode TFTs and DC-coupled measurement. C-D) Pixel circuit and corresponding optical image for enhancement-mode TFTs – necessitating pull-down switch architecture and pseudo-resistor with coupling capacitor to set DC bias. Red inset in B and D enclose a single pixel.

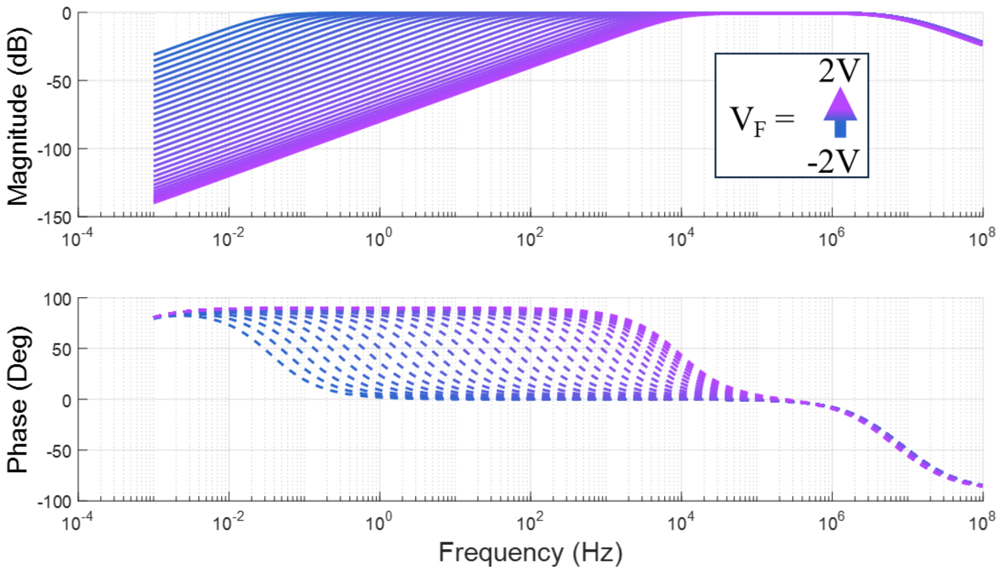


Figure 9 Simulated bandwidth for various pseudo-resistor biasing showing a per-pixel tunable high-pass filter.

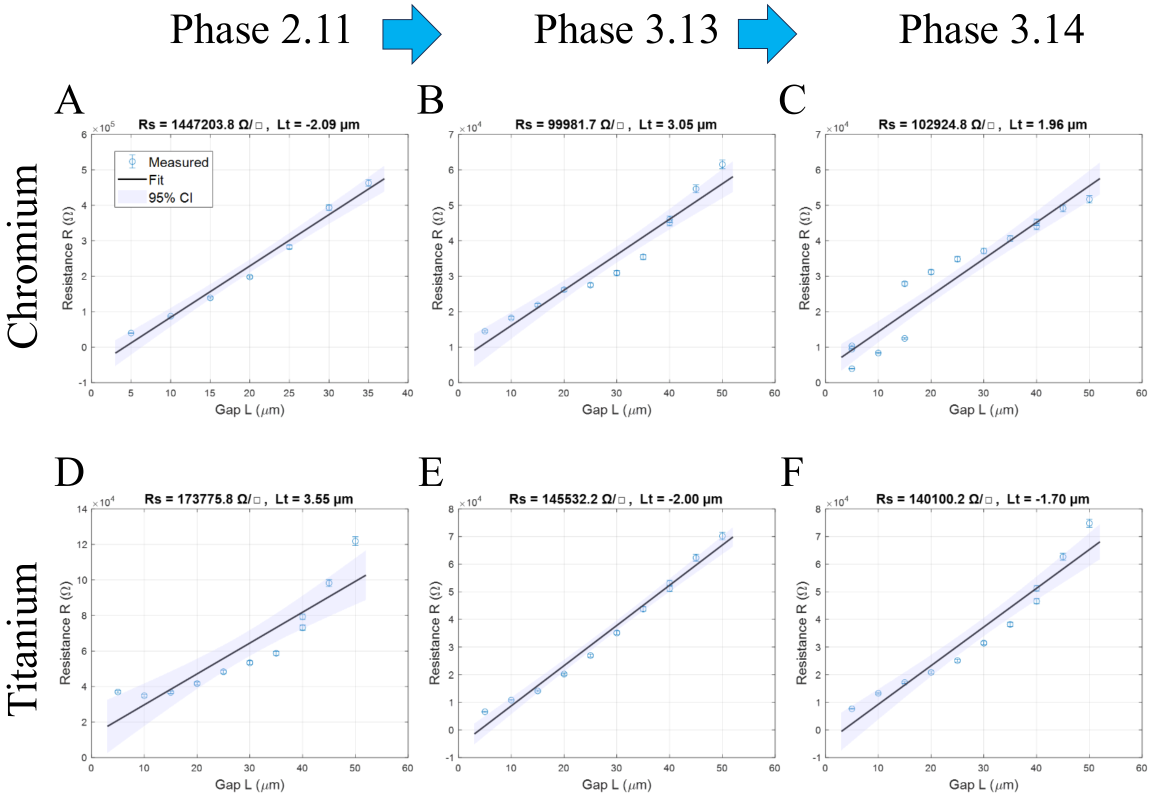


Figure 10 A four-point transmission line method was used to extract sheet resistance and contact resistance from an IGZO bar fabricated on test coupon patterns. A ±1uA sweep of current was passed through two outer contacts while the voltage drop across two inner contacts was measured for varying inter-contact edge-to-edge separation. A linear regression extracted the slope and intercept and a calculated 95% confidence interval is shown in light blue. A-C)

Chrome test coupon measurements across different phases of fabrication (naming convention shown in Supplementary Figure 1). D-E) Titanium test coupon measurements across different phases of fabrication.

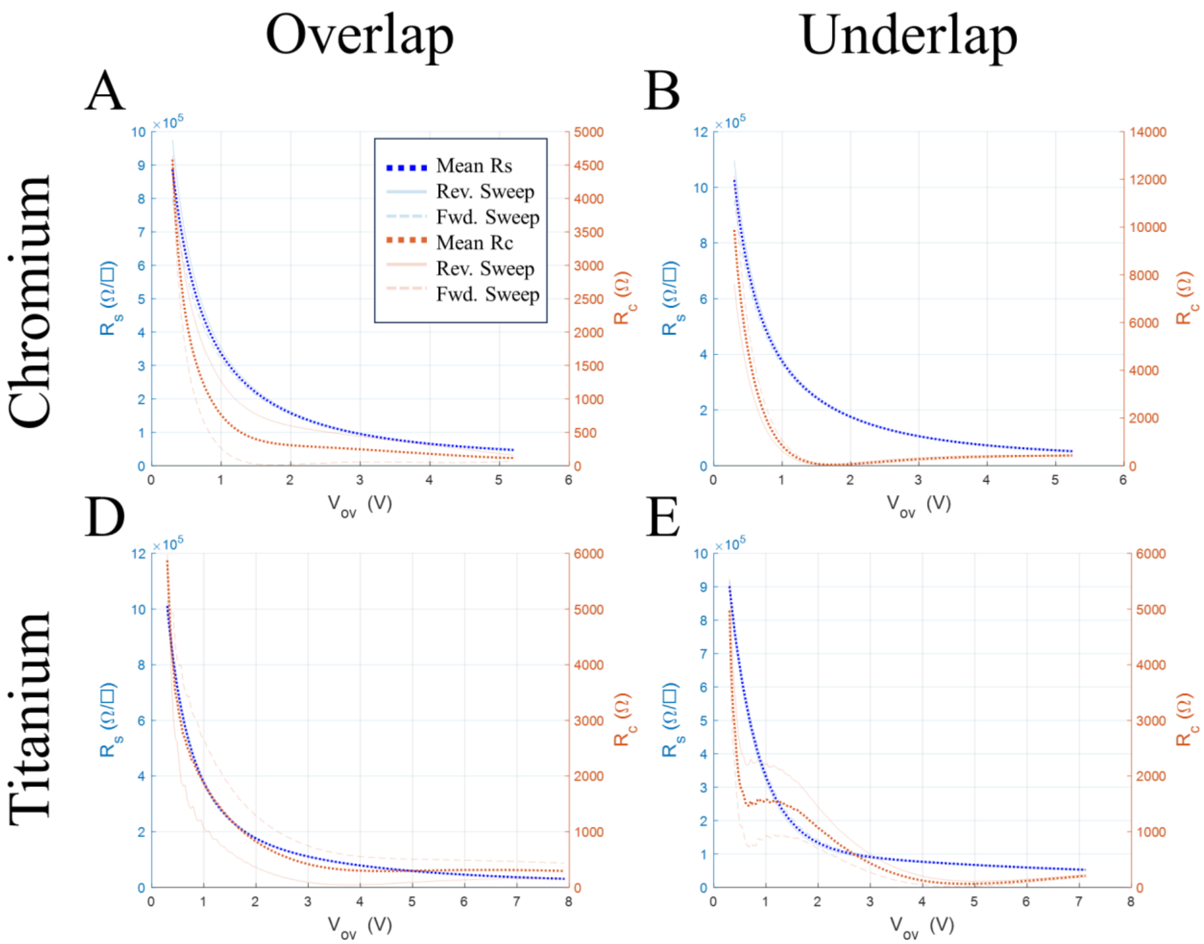


Figure 11 Gated transmission line model showing the effects of bottom-gate bias on the contact properties. All four panels show regression fits for sheet resistance and contact resistance as a function of overdrive voltage. The threshold voltage was independently extracted from the low-Vds transfer curves for all measured devices beforehand, and transfer curves with forward and reverse sweeps were used to estimate a range of values for fitted sheet and contact resistances. A-B) Compares the gated-TLM results between TFTs with and without overlapping gate-to-source and gate-to-drain regions for chromium, and C-D compares the same patterns for the titanium contact patterns.

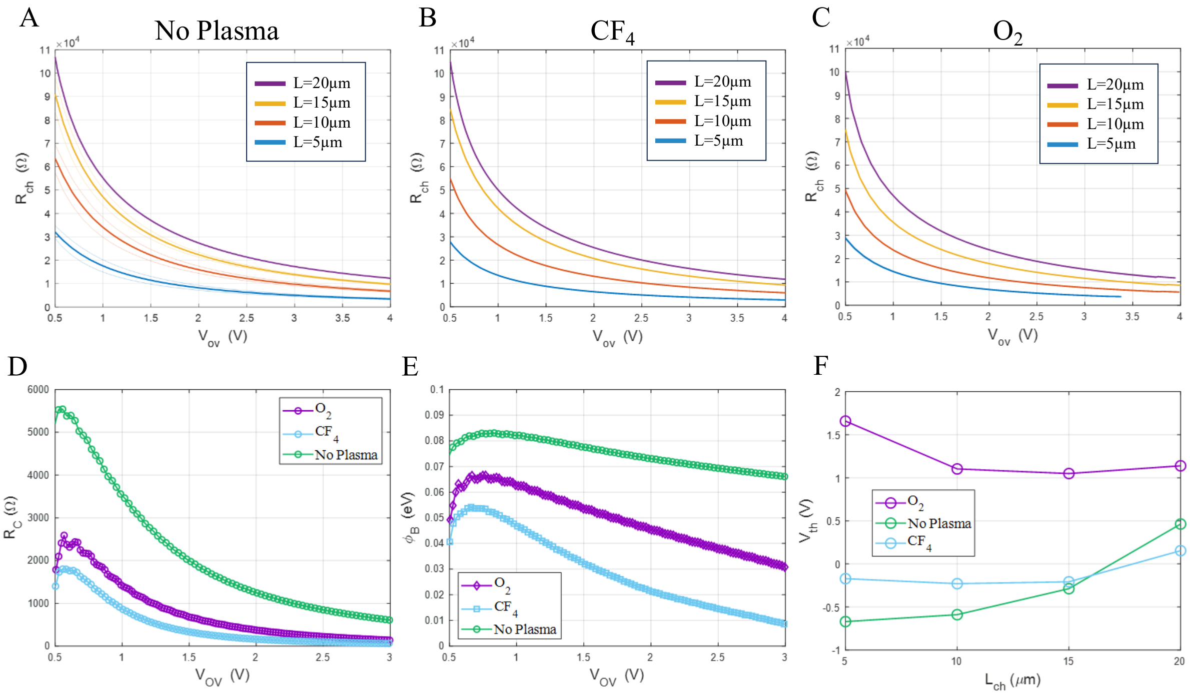


Figure 12 Plasma treatment study comparing different surface preparations prior to source and drain metallization. A-C) Total resistance plotted versus overdrive voltage for each channel length. D) Fitted contact resistance to the corresponding data in panels A-C. E) Extracted estimated Schottky barrier height. F) Threshold voltage extracted from low-Vds transfer curves for the corresponding data plotted in panels A-C.

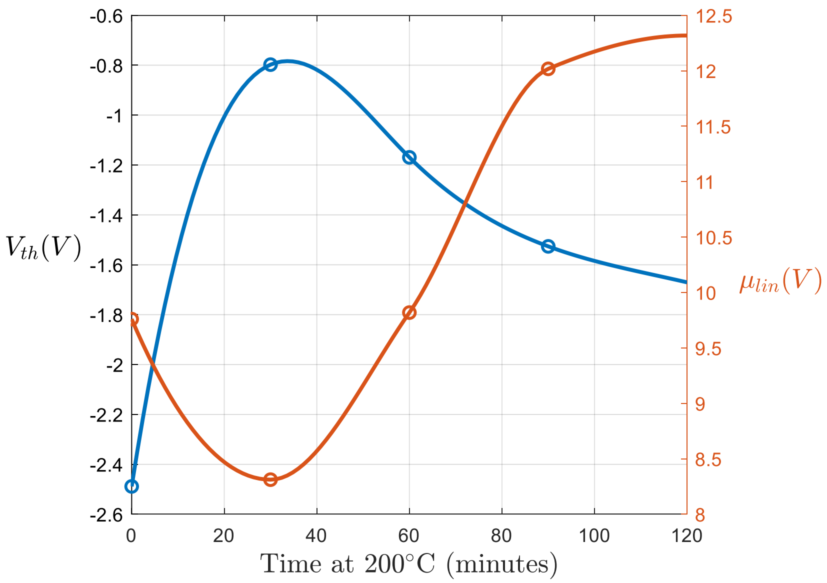


Figure 13 Extracted threshold voltage and linear mobility plotted as a function of annealing time for a depletion-mode dual-gate TFT with W/L of 356µm/10µm employing a chrome contact metal together with $CF_4$ plasma treatment.

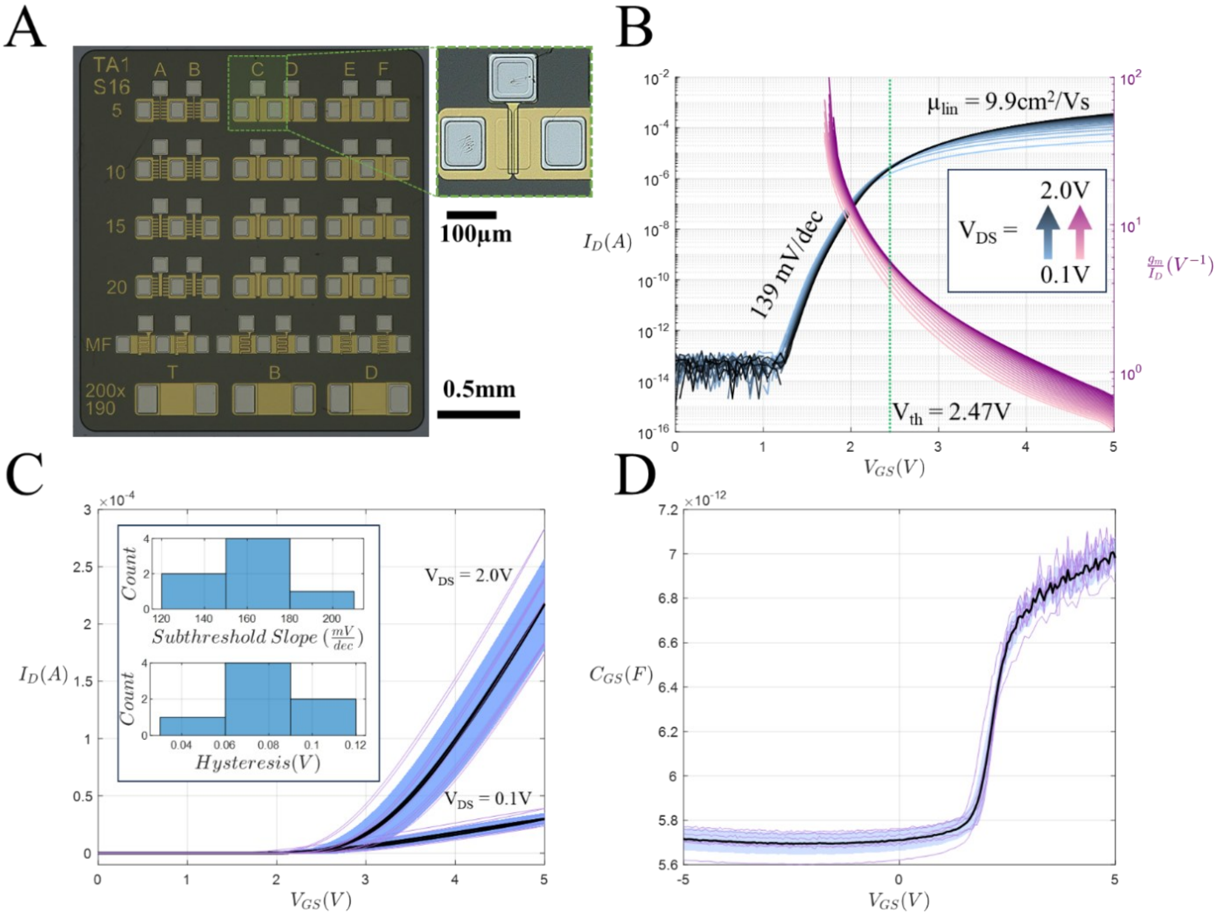


Figure 14 Overview of rectangular dual-gate TFT (device code C5) and corresponding characterization after implementing source/drain optimizations.

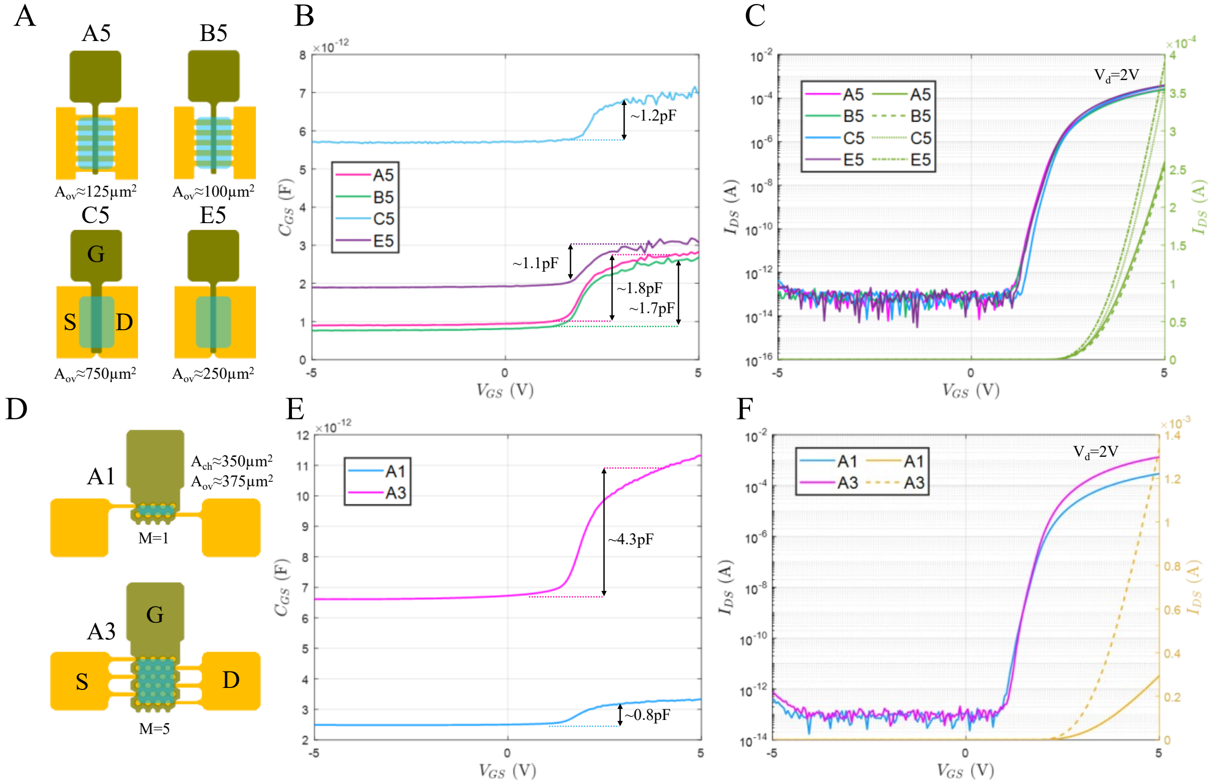


Figure 15 Various different layouts with corresponding capacitance-voltage and transfer curve measurements. A-C) Device characteristics for 100/5µm and varying overlap. D-F) Honeycomb device characteristics for single-finger (70/5µm) and 5-finger devices. Note that A3 corresponds to the HCA3 device code presented in Figure 2 P,Q in the main text.

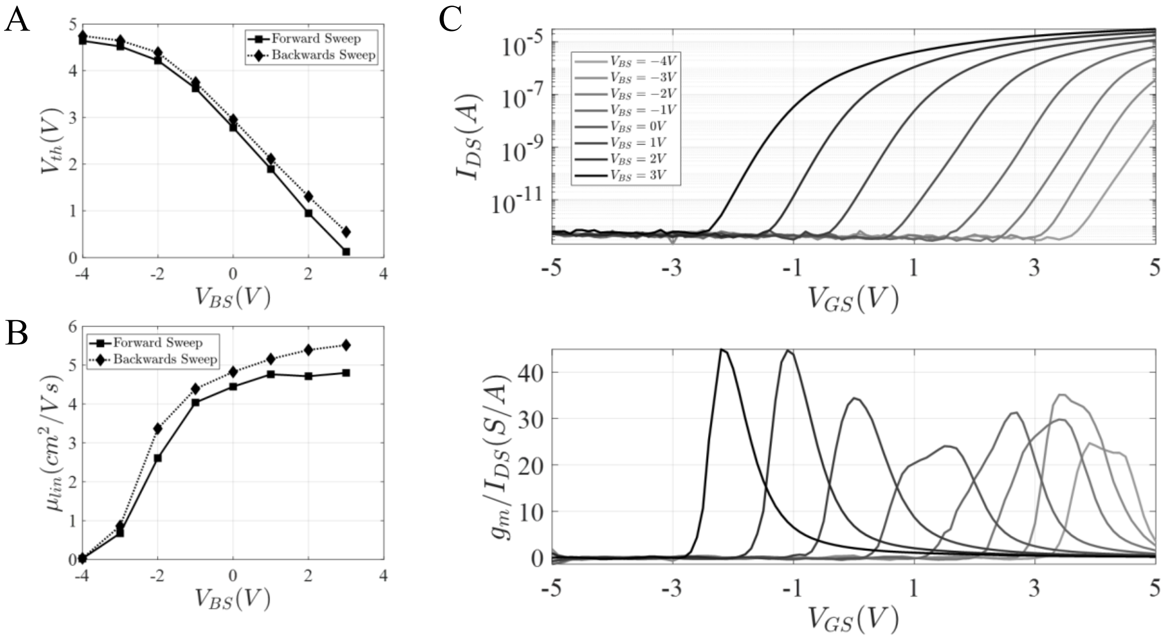


Figure 16 Transfer curves and corresponding extracted threshold voltage and mobility for split-gate architecture across different back-gate voltages.

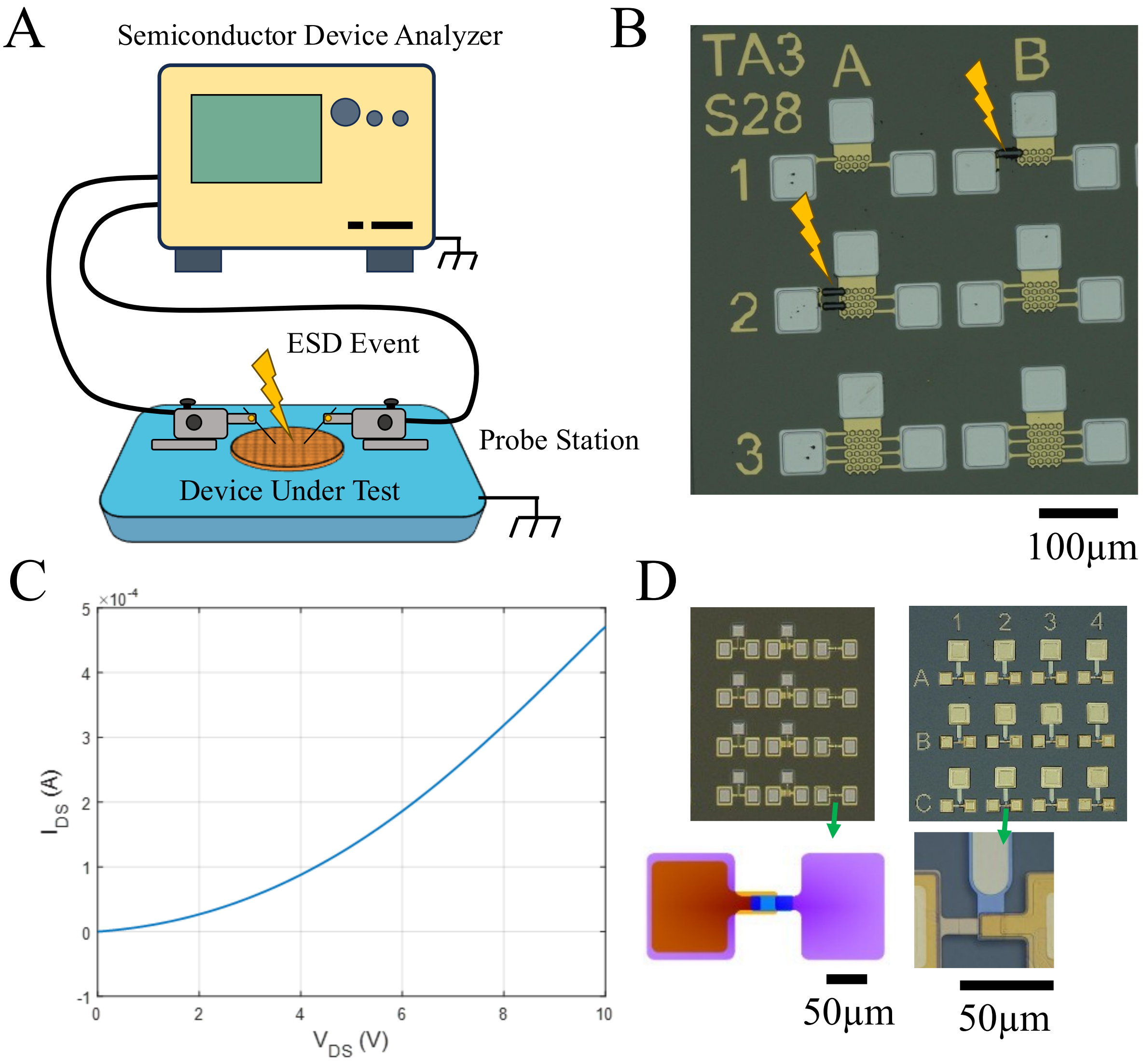


Figure 17 Representative figure showing ESD hazards for an insulated sample. Grounded probe stations still leave the devices under test floating when insulating polyimide substrates are used. Representative two-terminal and three-terminal diode-connected low-Vth TFTs are presented in panels C-D demonstrating rectifying behavior.

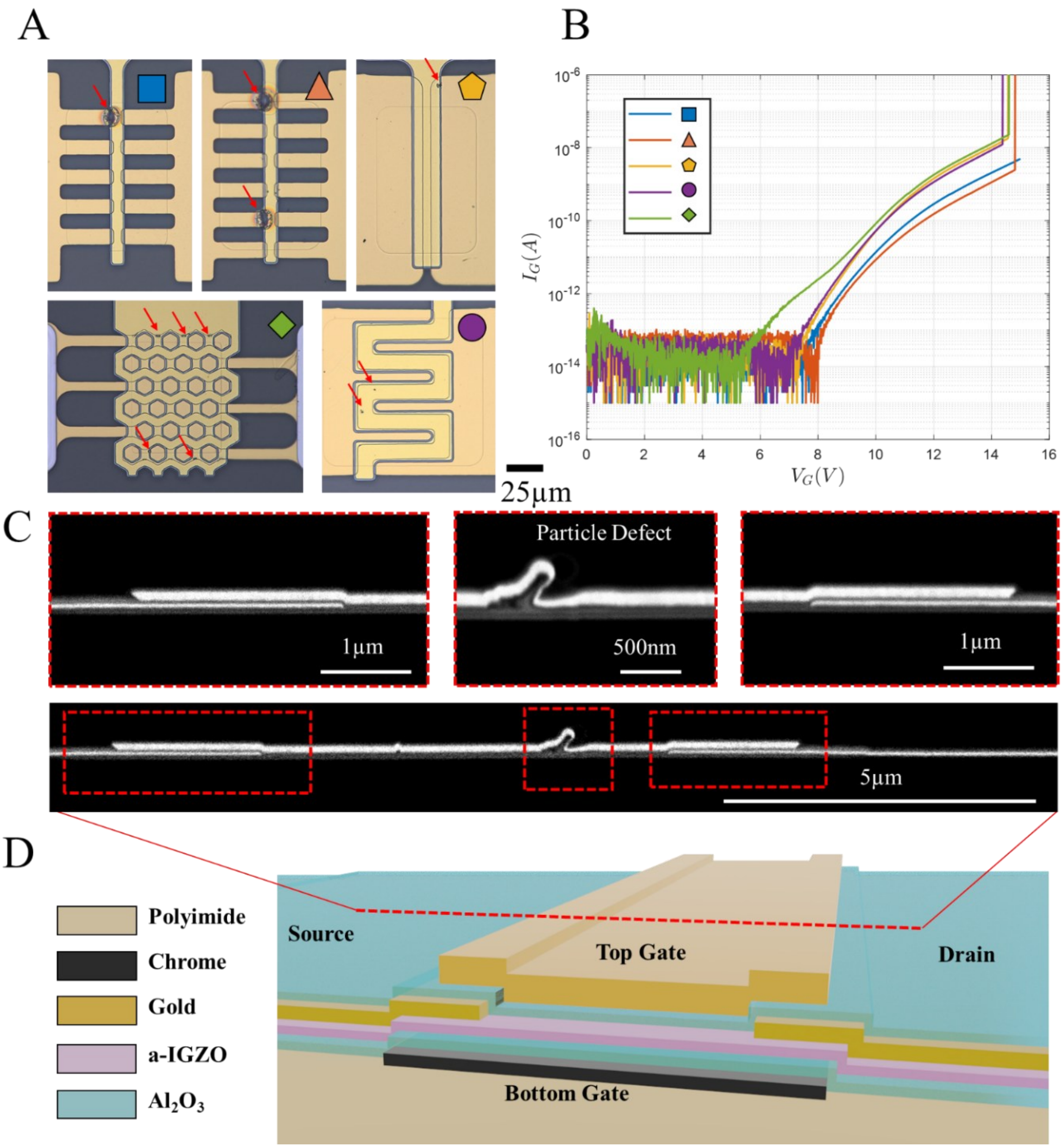


Figure 18 Overview of breakdown characteristics for five device architectures with source and drain grounded and gate voltage swept until dielectric breakdown was observed. A representative cross-sectional image is shown of a particle defect which can act as a source for premature breakdown.

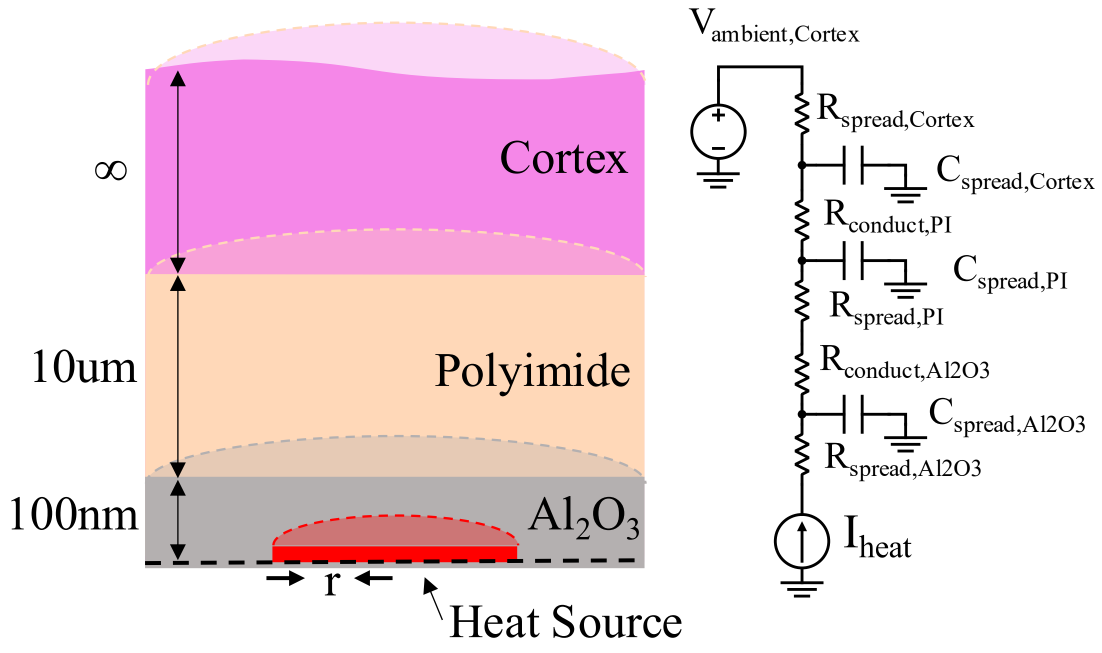


Figure 19 Cross-sectional view of cylindrically axis-symmetric implant model with corresponding electro-thermal heat dissipation model.

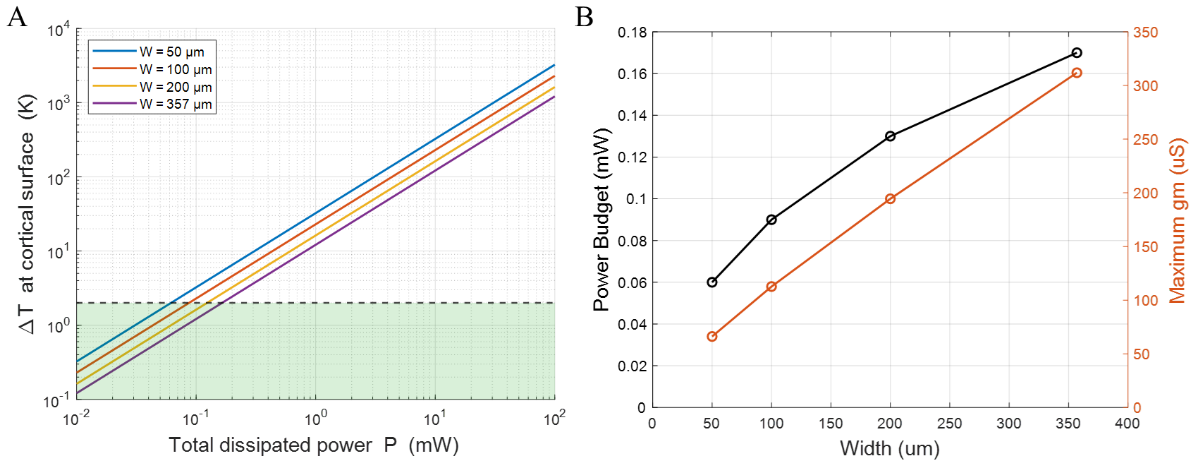


Figure 20 Simulated temperature elevation at the cortical surface and the resulting power budget for various channel widths (all with a fixed 5µm length). Maximum transconductances are extracted from measured transfer curves at the corresponding power budgets and appropriately scaled by W/L ratios.

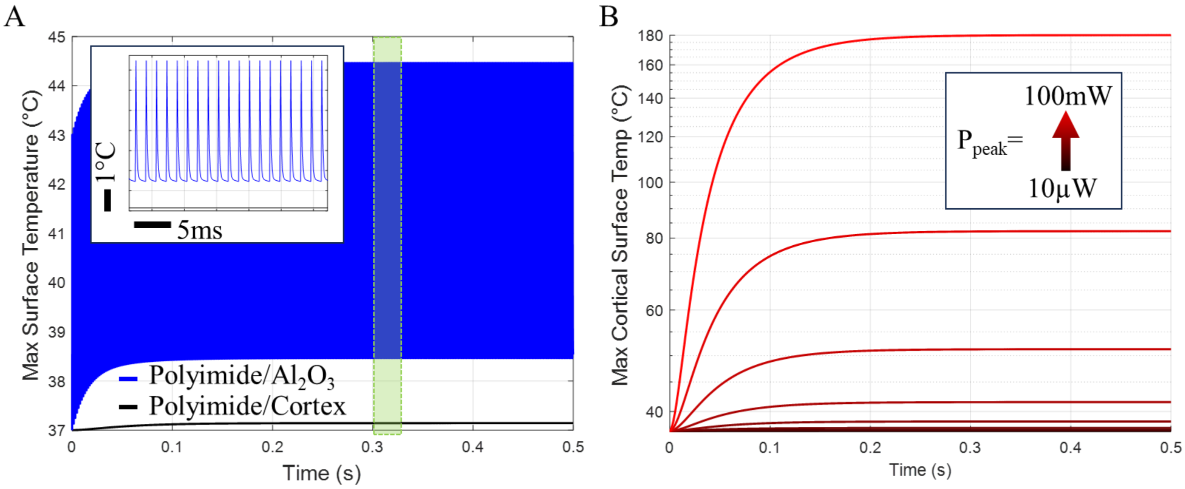


Figure 21 Transient electro-thermal simulation for representative 100x5µm device. A) Iamp = 100µW with a 100µs pulse period and 1.6ms period (1/16th duty cycling). B) Repeated simulations from A but with Iamp swept from 10µW to 100mW.

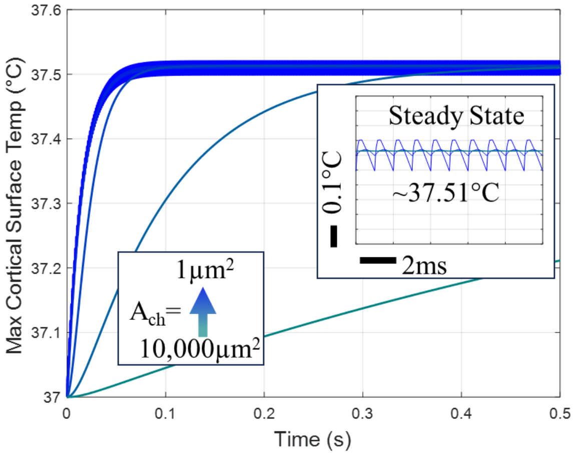


Figure 22 Transient electro-thermal modeling for duty-cycle scaling in lockstep with channel area scaling.

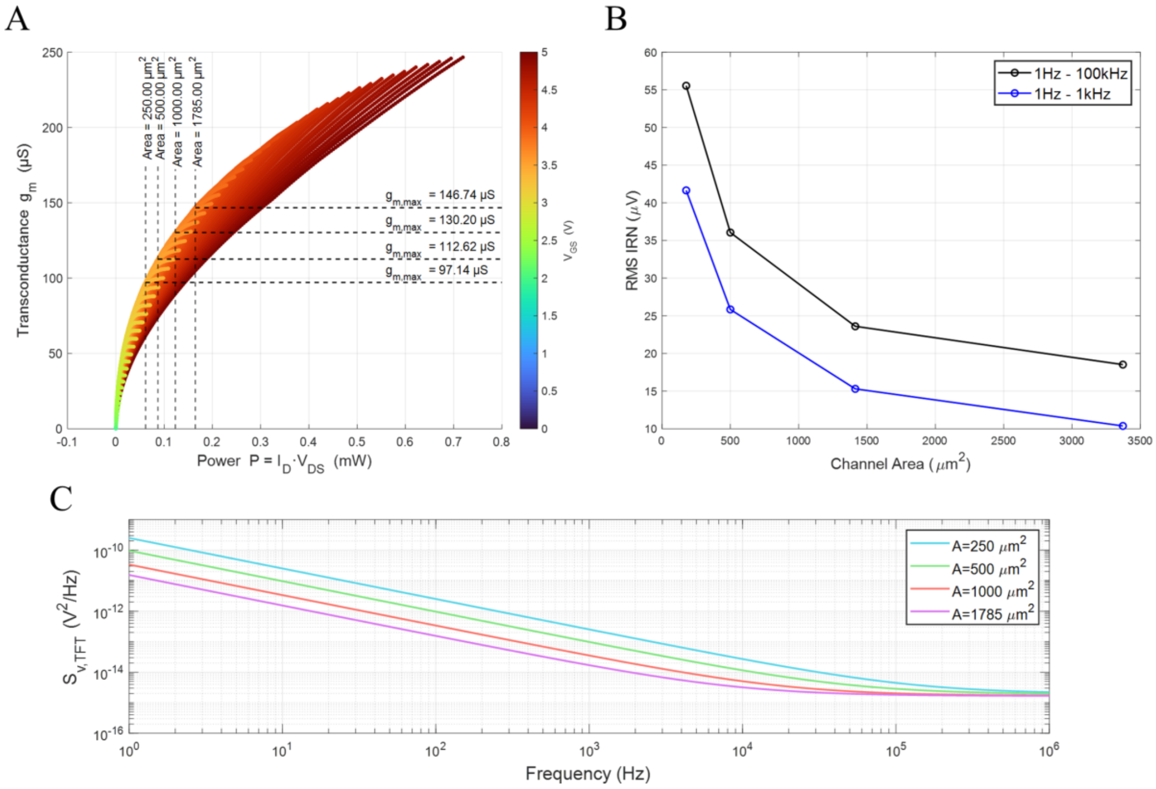


Figure 23 Extracted biasing from transfer curves associated with a W/L ratio of 20. A) Plotted transconductance as a function of bias-point and power with peak transconductance values shown for four representative channel areas and their respective maximum 2°C power budget. B) Total integrated RMS noise voltage referred to the gate for two representative bandwidths. Values correspond to maximum transconductance found in panel A. C) Input-referred noise voltage density calculated for four representative channel areas at the bias points determined in panel A.

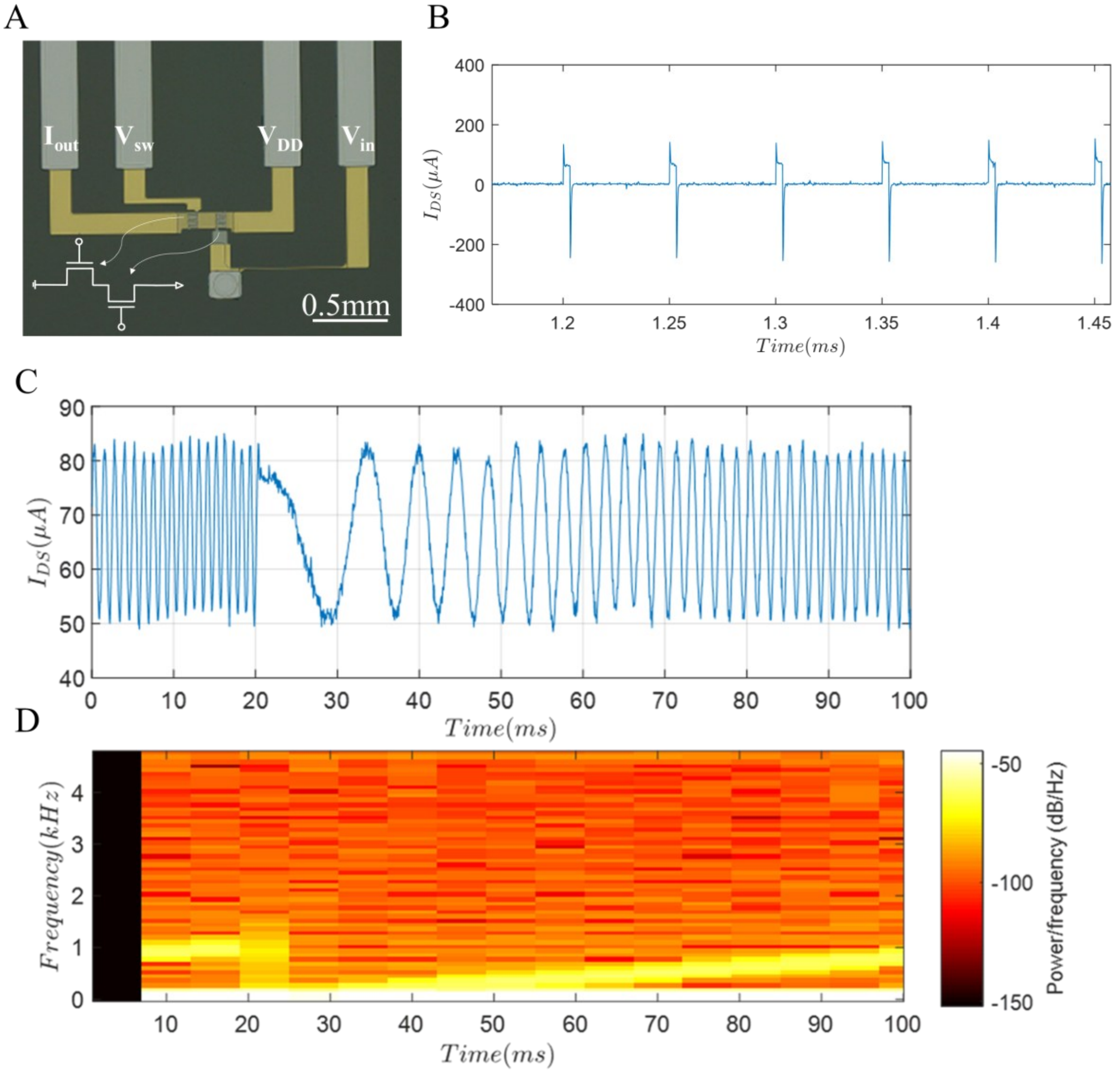


Figure 24 Benchtop measurement of single depletion-mode pixel. A) Optical image of test pixel composed of two 356µm x 10µm depletion-mode TFTs with corresponding schematic diagram. B) Representative window of current measured using a benchtop transimpedance amplifier and oscilloscope while $V_{sw}$ was driven with a square wave pulse with 6.25% duty cycle at 20kHz and $V_{in}$ was driven with a chirp signal. C-D) Demultiplexed waveform in time domain and a corresponding spectrogram showing the recovered chirp waveform with high SNR.

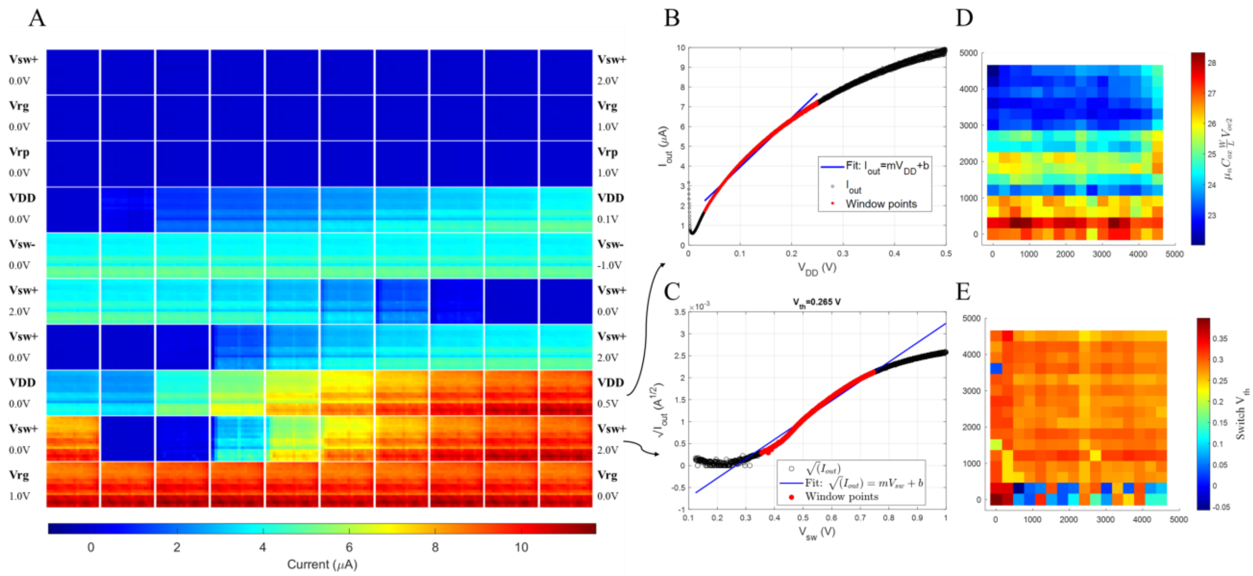


Figure 25 Overview of calibration sequence used to assess array performance. A) Tile plot with rows corresponding to a bias voltage sweep, and columns representing a linear sweeping between the left and right end-point voltages. The entire calibration sequence was carried out over a 10 second period. B-C) Representative curves during the indicated sweeps and corresponding fitted linear models used to extract key pixel parameters. D-E) Corresponding plots for the extractions shown in B and C.

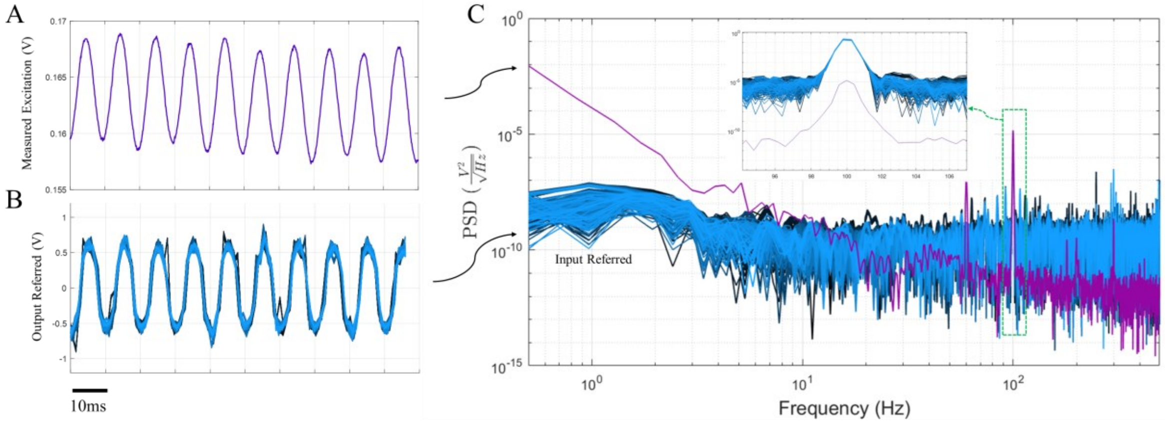


Figure 26 Benchtop saline measurements of injected 5mV amplitude 100Hz waveform for A) passive platinum microelectrode measured through benchtop oscilloscope and B) all 256-channel pixels for an array using the pseudo-resistor pixel architecture. C) Power spectral density for input-referred signals, scaled to match 100Hz signal power measured on passive microelectrode (pre-scaling shown in inset).

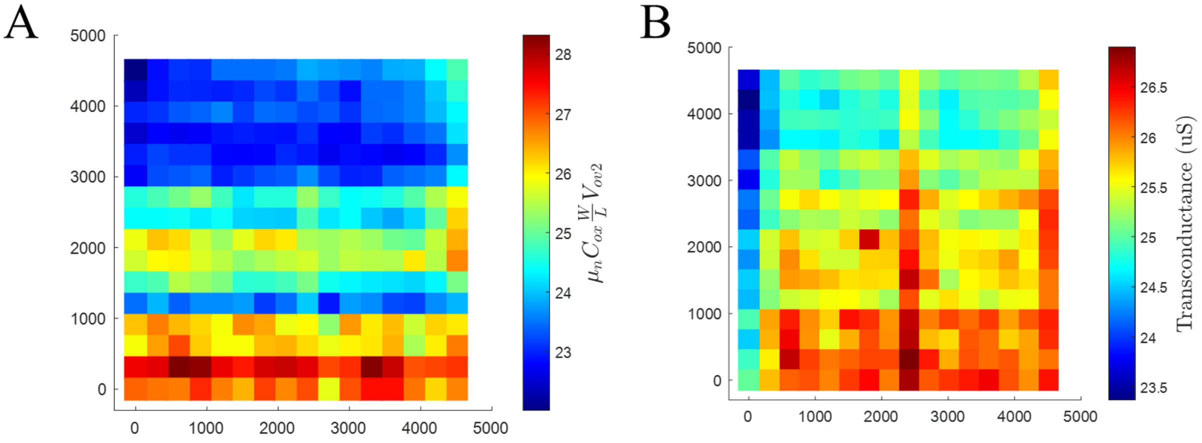


Figure 27 Comparison between pixel transconductance estimated through the calibration regression fitting method and the direct signal-injection method.

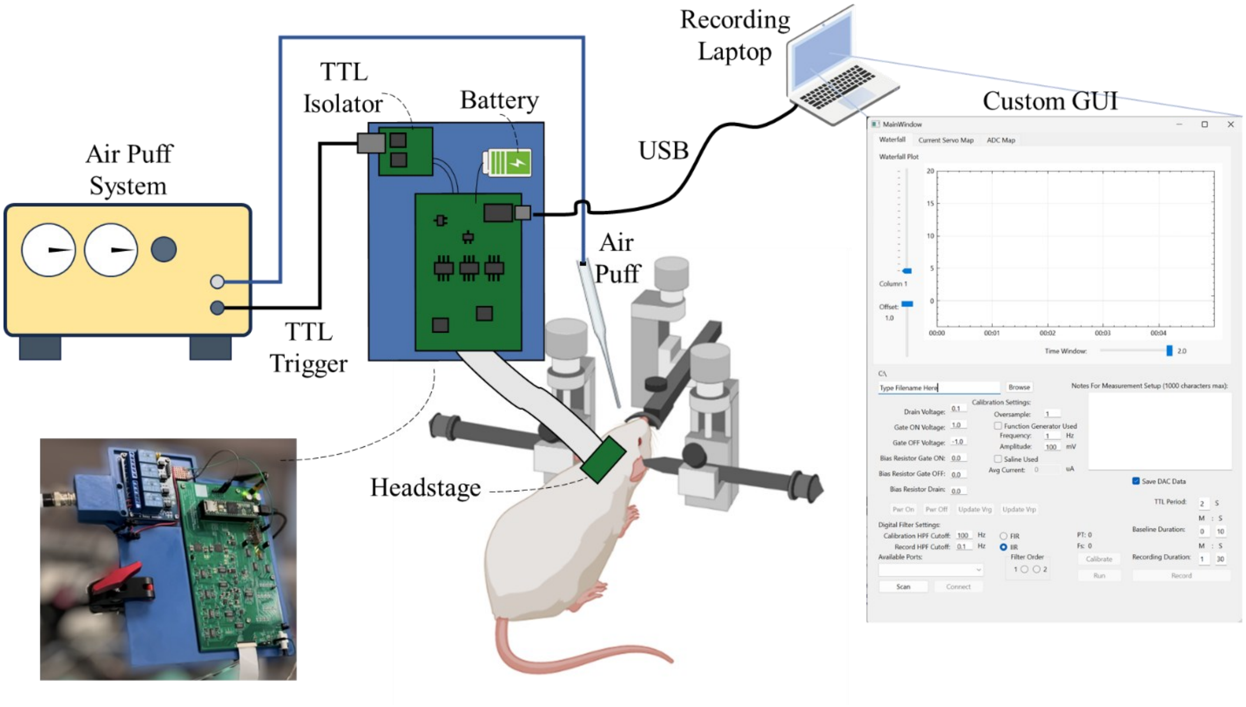


Figure 28 Overview of whisker air-puff stimulation experiment showing the custom external acquisition board connected to the TFT-based active μECoG array through a ribbon cable and headstage.

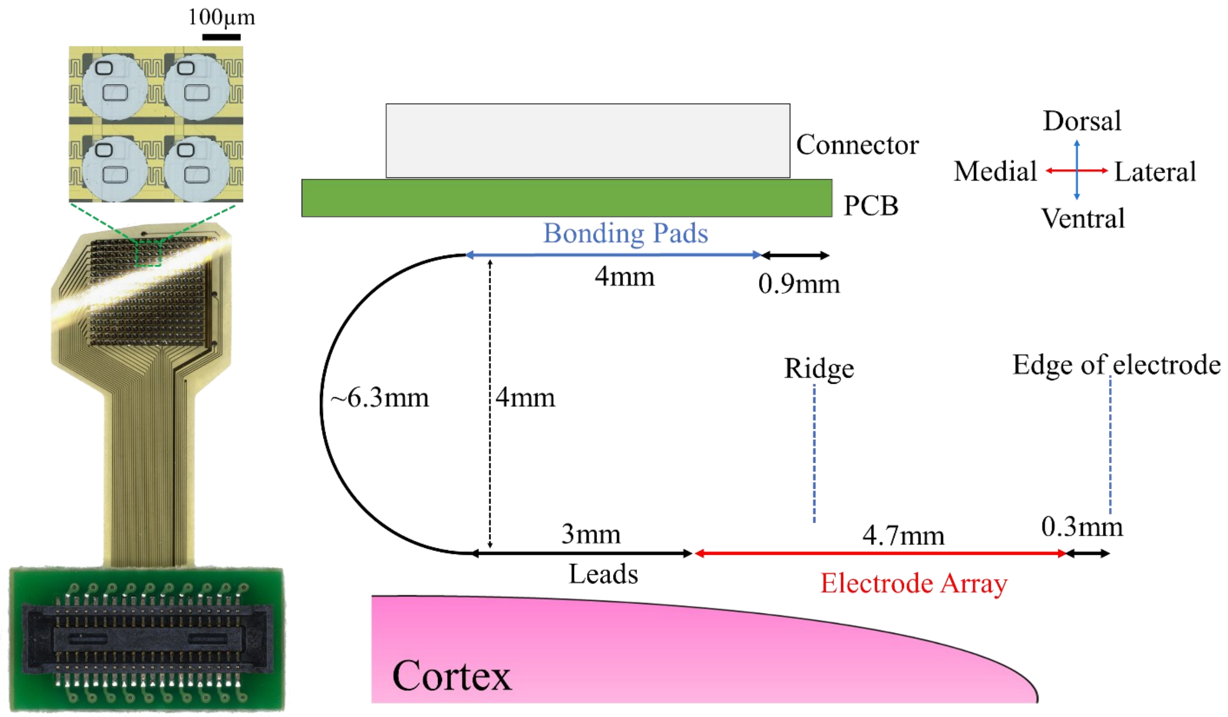


Figure 29 Design details for implanted active µECoG array showing a side-view of the chronic implant along with relevant sizing specifications.

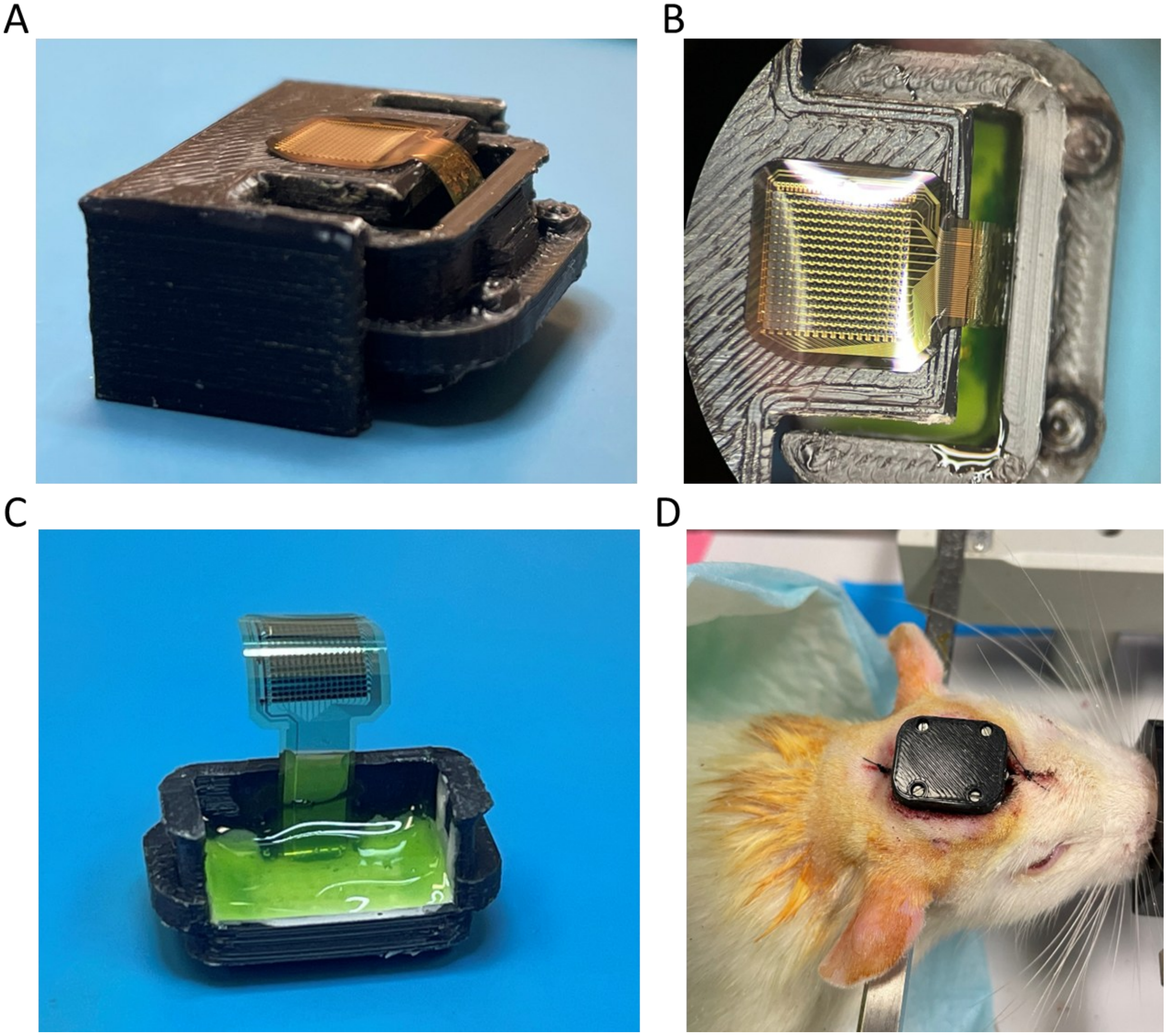

Figure 30 Overview of packaging and chronic surgical implantation. A-B) Sideview and top view of a 256-channel TFT-based ECoG array temporarily secured to an assembly jig used during packaging to achieve a fixed curvature profile and distance from the enclosure. C) Packaged array after material curing (with jig removed). D) Packaged array after implantation in Sprague-Dawley rat. A lid was secured to the packaged array shown in panel C to prevent damage to the connector.

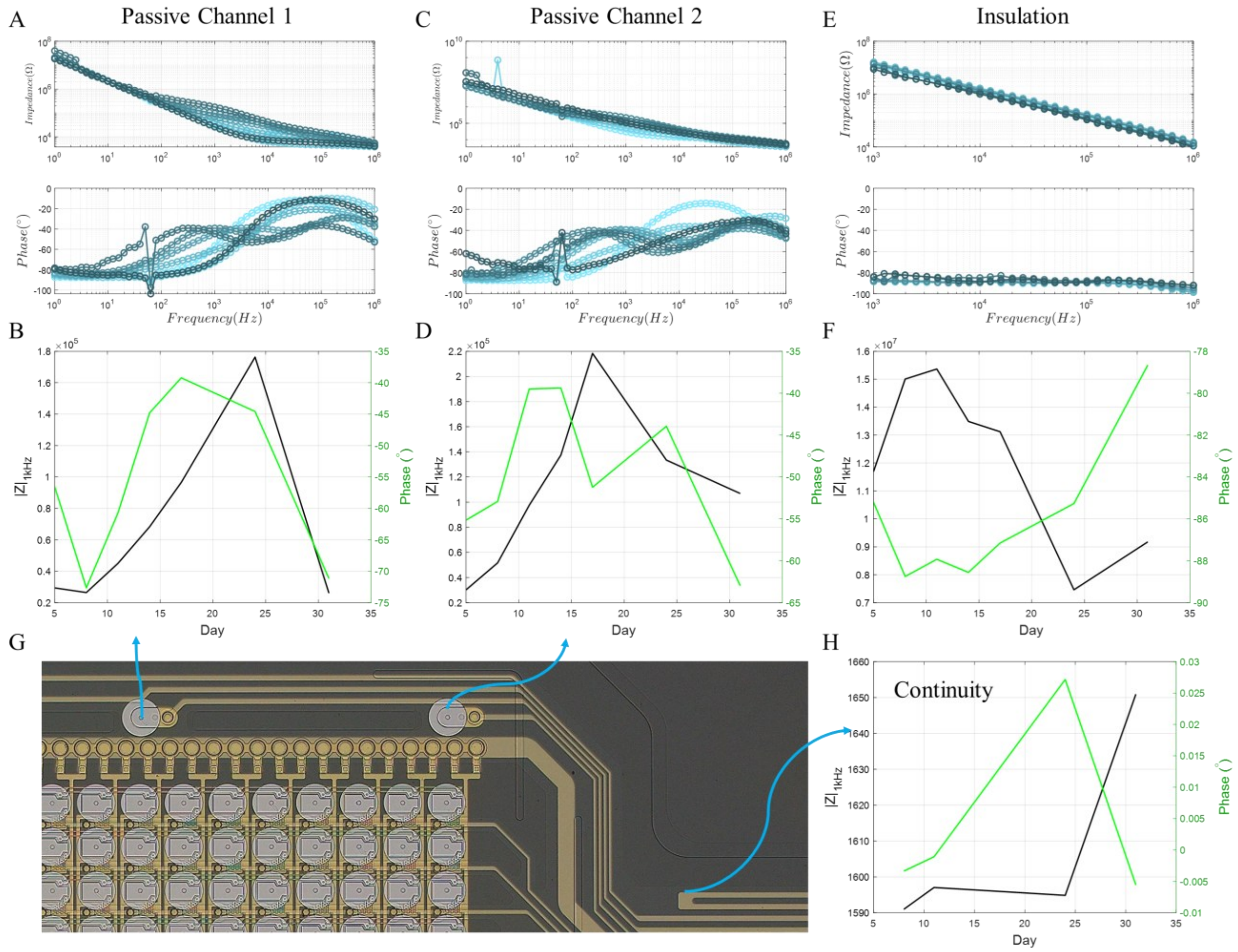


Figure 31 Overview of implanted passive platinum ECoG impedance over the duration of the chronic study. A-B) full impedance spectra of passive channel 1 with corresponding 1khz impedance magnitude and phase versus implant day. C-D) Same data as A-B for the second passive channel (both shown in panel G). E-F, H) Impedance spectra and corresponding 1khz impedance of a fully-passivated metal trace shown in panel G (blue arrow indication) along with the continuity between the two traces shown, indicating that the traces remained continuous (no open-circuits) and the insulation remained capacitive (no shorts to tissue). This data was expected based on accelerated aging studies, but was used as a safety metric that was monitored to ensure insulation quality during the chronic study.

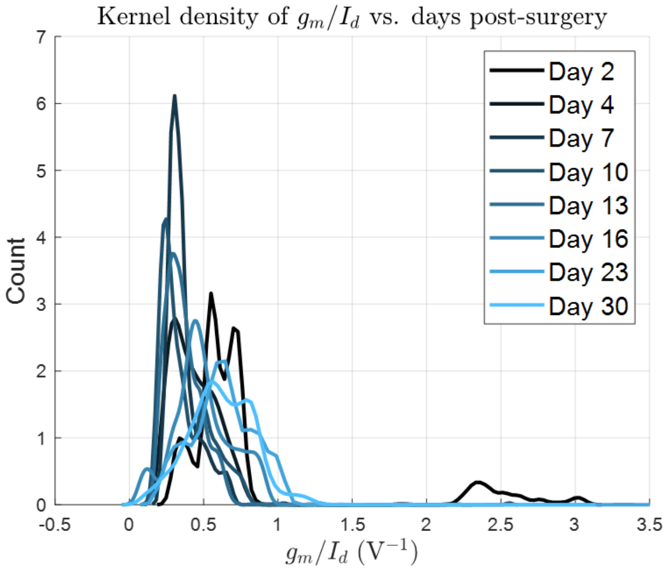


Figure 32 Summary histogram plot showing the normalized kernel density of extracted pixel sensing TFT transconductance normalized by the on-current. Distributions are calculated for all pixels across different recording files and biasing conditions.

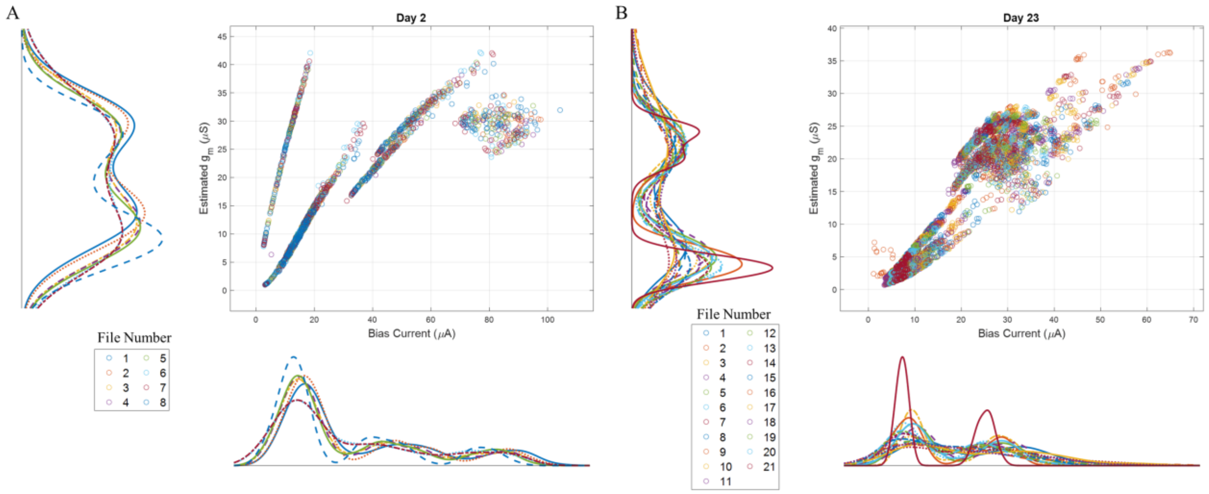


Figure 33 Two representative sampled bivariate histogram density plots for day two and twenty-three of the chronic implant study showing the correlation between estimated transconductance and bias current. Data is grouped into file numbers which correspond to a set of different calibration curves, and each plotted point represents a set of extracted values for a single pixel. Corresponding marginal kernel densities are plotted on their corresponding x and y axis.

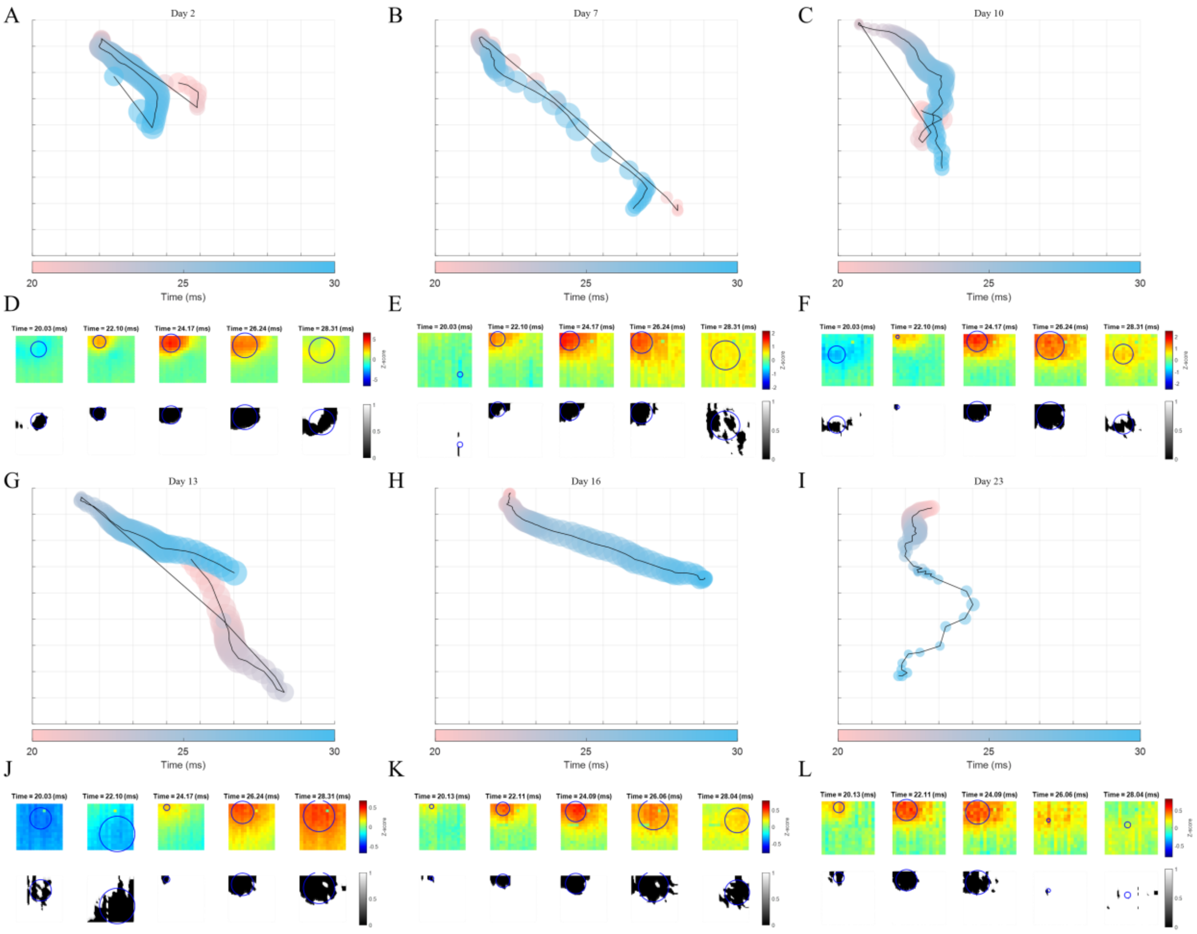


Figure 34 Weighted centroids of band-pass filtered trial averages from 20ms to 30ms after air-puff stimulation shown for representative recordings throughout the chronic study. Panels A-C and G-I show centroid trajectories with each marker area corresponding to the centroid radius, each marker color corresponding to the time, and each plot axis spanning the entire electrode array layout (0 to 4.5mm in both X and Y). Tile plots are shown for a subset of data in panels D-F and J-L with weighted centroids overlaid atop the z-score heatmap and each corresponding binary mask used to calculate the weighted centroid is plotted below.

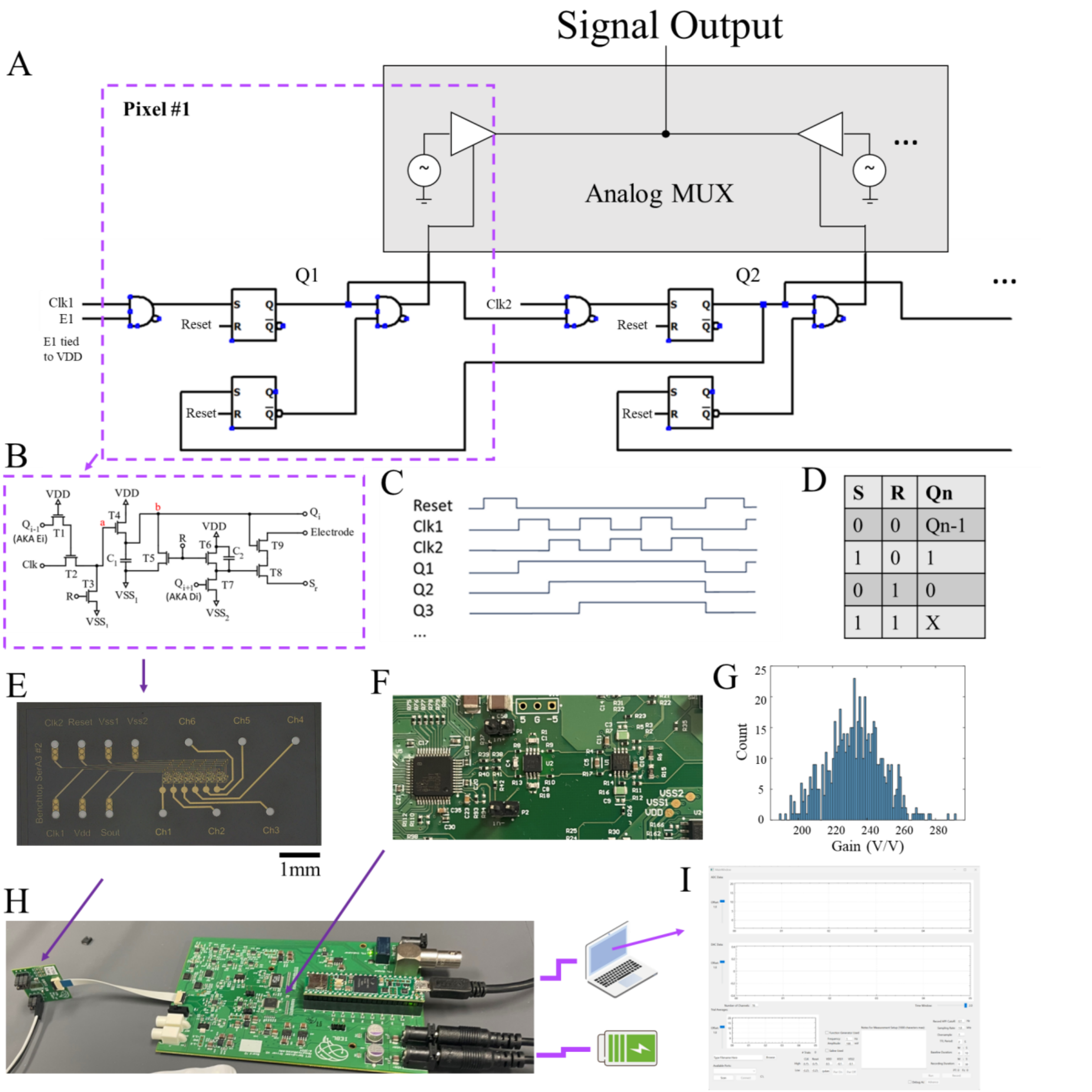


| S | R | Qn |
|---|---|---|
| 0 | 0 | Qn-1 |
| 1 | 0 | 1 |
| 0 | 1 | 0 |
| 1 | 1 | X |

Figure 35 Overview of first shift register architecture. A) Dynamic logic circuit diagram showing system concept for time-division multiplexed electrode array using "on-chip" gate driver circuits. A start enable signal, "E1" is input to the first cell (dotted purple box) and even and odd cells receive asynchronous clock inputs "CLK1" and "CLK2". B) cell circuit diagram showing the TFT-based dynamic logic circuit which utilizes a set-reset methodology along with pre-charging capacitors across all cells to known states. C) Timing diagram showing the propagation of data from cell 1 through the shift register. Note that the $i^{th}$ shift register cell output also acts to disable the $(i-1)^{th}$ cell, resulting in only one gate address being enabled at a given time. D) Truth table for each SR latch to help describe the operation. E) Optical die micrograph of a fabricated 6-channel shift register-based active microelectrode array. F) photograph of a custom acquisition system used to both drive and record from the circuit shown in panel E. G) Histogram of measured gain of the analog front end shown in panel F. The histogram was generated from sampled band-limited noise that was injected with a known bandwidth and integrated noise power, and the mean of the histogram represents the midband gain of the system which was designed to be ~230V/V. H-I) Full system overview showing the custom acquisition setup and daughter board which was used to connect the system to individual test circuits during benchtop testing.

Custom realtime acquisition software was developed to provide control of voltages and clock settings, to record, and to plot data in realtime.

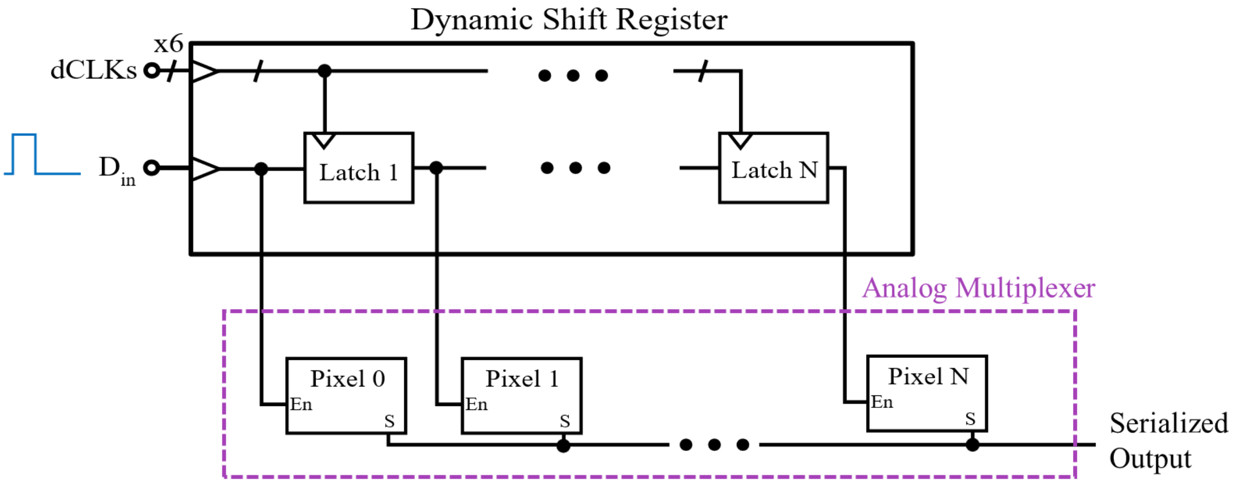


Figure 36 Diagram of second shift register based time-division multiplexing system which utilized different latch circuits and shift register architecture.

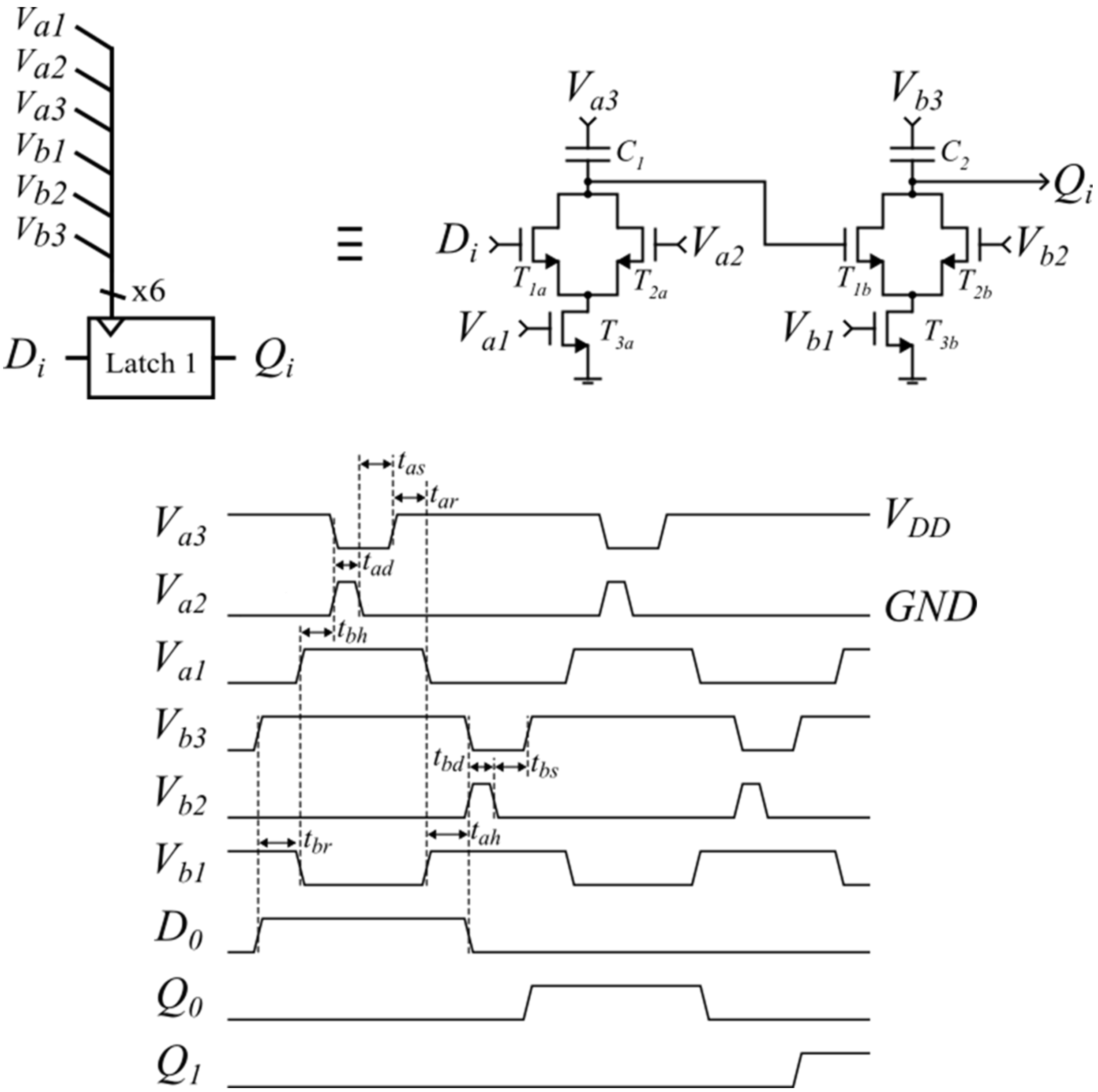


Figure 37 Additional details for second version of the dynamic shift register and corresponding timing diagram to demonstrate the concept. $t_{a,br}$: Duration for a and b sub-latches to read new state (i.e. charge C1,2). $t_{a,bh}$: Setup time for a,b sub-latch to hold. $t_{a,bd}$: Duration required to discharge a,b sub-latch capacitors. $t_{a,bs}$: Duration required for a,b sub-latch to settle before proceeding with read/transfer operations.

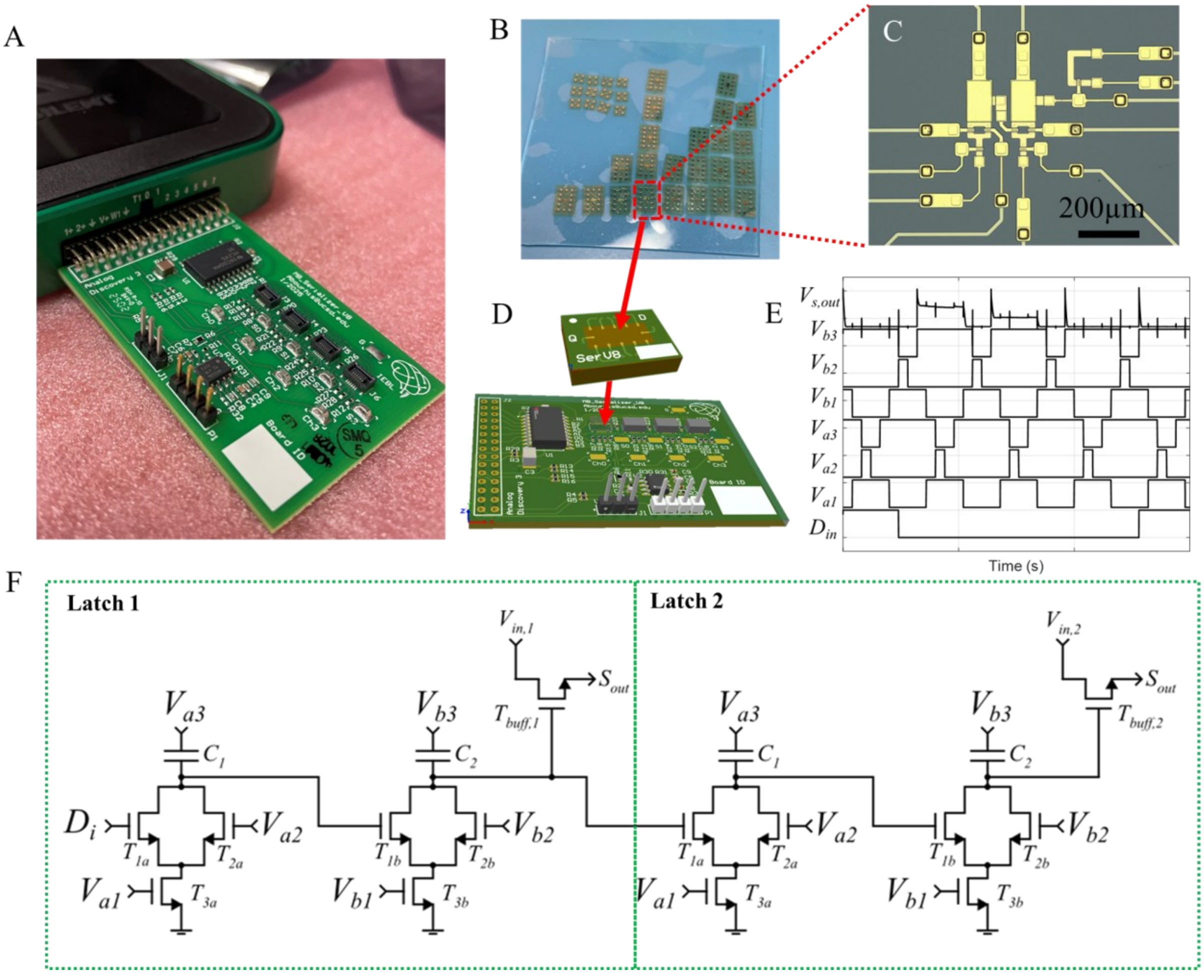


Figure 38 Overview of dynamic latch fabrication, assembly, and testing. A) Image showing custom motherboard plugged into a Digilent Analog Discovery 3. B-D) Photograph, optical die micrograph, and 3D CAD rendering of the flexible dynamic latches and daughter/mother boards used for benchtop testing. E) The time-division multiplexed output current, Sout, was fed to a transimpedance amplifier on the motherboard shown in A and D, and this amplifier was connected to a voltage measurement channel on the Digilent system, and recordings were taken in parallel to the digital clock recordings shown in panel E. Two test waveforms were applied at Vin,1 and Vin,2 shown in panel F. The output voltage shown in panel E demonstrates the system's ability to transfer the digital pulse applied to Di through the shift register, noting that Din in panel E is on for one clock period, then off for three, and since there are only two latches here, we see two output pulses followed by two zeros.

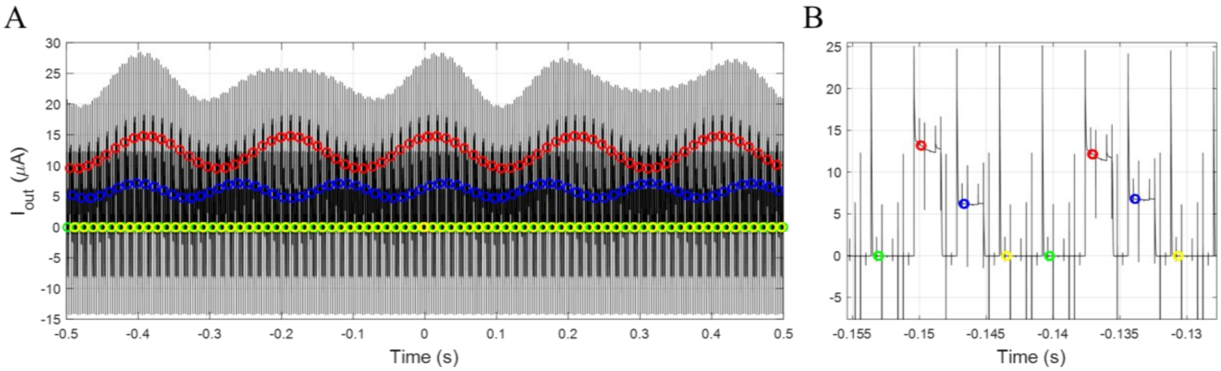


Figure 39 Demultiplexing the data from Supplementary Figure 31, showing the recovered output buffered waveforms which were injected into $V_{in,1}$ and $V_{in,2}$. Two distinct frequencies were injected to assess cross-talk caused by faulty state-transfer between latches, and as shown in panel A, clean sinusoids were recovered without any observed inter-channel mixing.

### Supplementary Tables

| **Material** | **Thermal Processing**† | **εr (1 kHz-1 MHz)** | **Volume ρ (Ω cm)** | **WVTR*** **($g\ m^{-2}\ day^{-1}$)** | **Young's E (GPa)** |
|---|---|---|---|---|---|
| Bulk Ti (Grade 2/5) [65–67] | Laser welding < 400 °C | metal (∞) | $4.2x10^{-5}$ | undetectable‡ | 110 |
| Bulk $Al_2O_3$ [68,69] | Sintering 1500-1600 °C | 9.0-9.8 | > $10^{14}$ | undetectable‡ | 300-380 |
| Bulk Borosilicate Glass [70] | Glass-to-metal seal 450-600 °C | 4.6 | $10^{14}$-$10^{15}$ | undetectable‡ | 64 |
| Parylene C [71–73] | CVD 25-40 °C | 3.1 | $8.8 \times 10^{16}$ | 1.6-16 (50-5 μm) | 2.5-3.2 |
| Polyimide (PI-2600) [74-76] | Oven Curing 350 °C | 3.1-3.3 | ≥ 10^16 | 2-20 (50-5 μm) | 3-8.5 |
| PDMS (Sylgard 184) [77-79] | Curing 25°C-150 °C | 2.5-2.7 | $1x10^{14}$ | 24-430 (50-100 μm) | $1.8x10^{-3}$ |
| LCP (Vectra A950 / R-flex 3600) [80,81] | Melt-laminate 280-320 °C | 2.9-3.2 | $5x10^{15}$ | 0.3 (25 μm) | 2-4.8 |
| ALD $Al_2O_3$ [82,83] | ALD 80-300 °C | 8-10 | > $10^{14}$ | ≤ $1.7x10^{-5}$ (25-50 nm) | 120-200 |
| ALD $HfO_2$ [84,85] | ALD 80-300 °C | 20-25 | > $10^{12}$ | $10^{-5}$-$10^{-6}$ (10-50nm) | 240-370 |
| ALD $Al_2O_3$ + Parylene [86] | ALD + CVD ≤ 200 °C | effective ≈ 3-8 | > $10^{14}$ | < $10^{-6}$ (20-30 nm) | 10-50 |

Table 1 Summary of typical encapsulation materials and their relevant material and processing properties. †Rough estimates for peak substrate temperature during processing. *Numbers are highly dependent on thickness, temperature, and relative humidity. ‡Based on measurement limitations for ASTM F1249, ASTM F3299, and ISO 15106-3 standards.

| Compound | ΔfG°, 298 K(kJ mol-¹) | Number of Oxygen Atoms | ΔG°/O (kJ mol-¹ O) |
|---|---|---|---|
| $In_2O_3$ | – 827.23 | 3 | – 275.74 |
| $Ga_2O_3$ | – 998.3 | 3 | – 332.77 |
| ZnO | – 320.50 | 1 | – 320.50 |
| $TiO_2$ (rutile) | – 889.52 | 2 | – 444.76 |
| $Cr_2O_3$ | – 1053.11 | 3 | – 351.04 |

Table 2 Summary of binary oxides and their corresponding enthalpy of formation.[46-49]

| Layer | Thickness $t_i$ (µm) | Thermal Conductivity $k_i$ ($Wm^{-1}K^{-1}$) | Density $\rho_i$ (kg $m^{-3}$) | Specific Heat $c_i$ (J $kg^{-1}$ $K^{-1}$) | $R_{spread,i}$ ($KW^{-1}$) | $R_{conduct,i}$ ($KW^{-1}$) | $C_{spread,i}$ ($JK^{-1}$) |
|---|---|---|---|---|---|---|---|
| $Al_2O_3$ (ALD)[33,55,56] | 0.1 | ~1.0 – 3.0 | ~2770 | ~880 | 9.85 x $10^3$ | 100 | 1.10 x $10^{-9}$ |
| PI2611[57,58] | 10 | ~0.033 – 0.15 | ~1400 | ~1090 | 1.01 x $10^5$ | 2.19 x $10^5$ | 6.87 x $10^{-8}$ |
| Cortex [59,60] | - | ~0.5-0.6 | ~1040 | ~3600 | 2.29 x $10^4$ | - | 1.97 x $10^{-6}$ |

Table 1 Summary of modeling parameters for an exemplar device with 100x5 µm channel area.

| Date | Experiment | Electrode ID | Data Collected | Included in Manuscript |
|---|---|---|---|---|
| 4/24/2025 | Chronic Implant | V8S1A17 | Imaging, stimulation evoked responses, electrochemical impedance | Yes |
| 3/1/2025 | Chronic Implant Pilot | V8S1A11 | Imaging, faulty electrode data (ESD damage) | No |
| 11/11/2024 | Acute | V7S5A2 | Imaging, TFT electrode recording | No |
| 4/3/2024 | Acute | V6S4A6 | Imaging, passive electrode recording, faulty TFT electrode (damage during surgery) | No |
| 1/4/2024, 1/18/2024, 1/26/2024, 2/2/2024 | Chronic Pilot | V6S4A7 | Imaging, TFT electrode recording, electrochemical impedance | No |
| 1/10/2024 | Acute | V6S4A6 | Imaging, passive electrode recording, TFT electrode recording | Yes |

| | | | | |
|---|---|---|---|---|
| 12/15/2023 | Acute | V6S4A6 | Imaging, TFT electrode recording | No |
| 11/16/2023 | Acute | Not Recorded | Imaging, TFT electrode recording | No |
| 10/24/2023 | Acute + post-mortem chronic pilot | V6S4A2 | Imaging, passive electrode recording, TFT electrode recording | No |
| 9/26/2023 | Acute | V6S4A3 | Imaging, passive electrode recording, TFT electrode recording | No |
| 3/29/2023 | Acute | Not Recorded | Imaging, passive electrode recording, TFT electrode recording, FLIR temperature measurement, Histology | Yes |
| 2/17/2023 | Acute | Not Recorded | Imaging, passive electrode recording, TFT electrode recording | No |
| 11/3/2023, 10/20/2023, 10/4/2023 | Chronic Pilot | Not Recorded | Imaging, passive electrode recording, TFT electrode recording | No |
| 10/3/2023 | Chronic Pilot | Passive Electrode | Imaging (testing housing designs) | No |

Table 2 Summary of animal recordings, including pilot studies used to help develop and iterate on electrode designs and surgical procedures.
